# Spinal cord injury reprograms muscle fibro-adipogenic progenitors to form heterotopic bones within muscles

**DOI:** 10.1101/2021.11.04.467192

**Authors:** Hsu-Wen Tseng, Dorothée Girard, Kylie A. Alexander, Susan M Millard, Frédéric Torossian, Adrienne Anginot, Whitney Fleming, Jules Gueguen, Marie-Emmanuelle Goriot, Denis Clay, Beulah Jose, Bianca Nowlan, Allison R. Pettit, Marjorie Salga, François Genêt, Marie-Caroline Le Bousse-Kerdilès, Sébastien Banzet, Jean-Pierre Levesque

## Abstract

The cells-of-origin of neurogenic heterotopic ossifications (NHO), which develop frequently in the periarticular muscles following spinal cord injuries (SCI) and traumatic brain injuries, remain unclear because the skeletal muscle harbors two progenitor cell populations: satellite cells (SCs) which are myogenic, and fibro-adipogenic progenitors (FAPs) which are mesenchymal. Lineage-tracing experiments using the Cre recombinase /LoxP system were performed in two mouse strains with the fluorescent protein ZsGreen specifically expressed in either SCs or FAPs in the skeletal muscles under the control of the *Pax7* or *Prrx1* gene promotors respectively. These experiments demonstrate that following a muscle injury, SCI causes the upregulation of PDGFRα on FAPs but not SCs and the failure of SCs to regenerate myofibers in the injured muscle, with instead reduced apoptosis and continued proliferation of muscle resident FAPs enabling their osteogenic differentiation into NHO. No cells expressing ZsGreen under the *Prrx1* promoter were detected in the blood after injury suggesting that the cells-of-origin of NHO are locally derived from the injured muscle. We validated these findings in the human pathology using human NHO biopsies. PDGFRα^+^ mesenchymal cells isolated from the muscle surrounding NHO biopsies could develop ectopic human bones when transplanted into immunocompromised mice whereas CD56^+^ myogenic cells had a much lower potential. Therefore, NHO is a pathology of the injured muscle in which SCI reprograms FAPs to uncontrolled proliferation and differentiation into osteoblasts.

## INTRODUCTION

Neurogenic heterotopic ossifications (NHO) are pathological heterotopic bones that develop in peri-articular muscles^1, 2^ following severe lesion of the central nervous system (CNS) such as spinal cord injuries (SCI), traumatic brain injuries, strokes and cerebral anoxia. NHO are frequent with an incidence of 10-23% in patients of traumatic brain injury, 10-53% in SCI patients, increasing to 68% in victims of severe combat blast injuries involving the spine^3–8^. NHO develop most frequently at the hip, elbow, knee and shoulder^2, 9^. Because of their large size, NHO are very incapacitating, causing significant pain and gradual reduction in the range of motion of affected limbs often progressing to complete ankylosis^2, 9^. This exacerbates functional disabilities by increasing difficulty in sitting, eating and dressing^10^. NHO growth can also cause nerve and blood vessel compression, further increasing patient morbidity^11, 12^. Despite knowing this pathology for just over 100 years, treatment is currently limited to surgical resection after NHO have matured^2, 10, 13, 14^, a procedure that is challenging, particularly when ossifications entrap the whole joint and adjacent large blood vessels and nerves. The development of improved treatments for NHO has been slow and trials of pharmacological interventions have continued to show limited effectiveness^15, 16^, reflecting the current limited knowledge on the etiology, pathogenesis and pathobiology of NHO. Indeed, the initial causal mechanisms that trigger NHO are unique to this pathology as they involve a severe CNS trauma^17, 18^ rather than other forms of trauma such as body burn, or activating mutations of osteogenic genes such as bone morphogenetic protein type I receptor *ACVR1* in fibrodysplasia ossificans progressiva (FOP)^17^. Therefore, it is essential to uncover the mechanisms of NHO pathogenesis to identify potential therapeutic targets and treatments that will reduce NHO development and remove the need for very invasive and delicate surgical resections^17^.

Our group previously reported the first NHO mouse model in which NHO spontaneously develop when a SCI is combined with a muscle injury without additional non-physiological manipulations such as the overexpression of bone morphogenetic protein (BMP) transgenes or the insertions of hyper-active mutants of BMP receptors^19^. Our model revealed that SCI causes a further exacerbation of the inflammatory response in injured muscles with exaggerated Ly6C^bright^ inflammatory monocyte / macrophage infiltration and persistent accumulation of the inflammatory cytokine oncostatin M (OSM), leading to persistent activation of JAK1/2 tyrosine kinases and signal transducer and activator of transduction-3 (STAT3) which in turn promote NHO instead of muscle repair^19–21^.

How injured skeletal muscles generate heterotopic bones instead of regenerating functional myofibers following severe lesions of the CNS remains a fascinating stem cell biology question and may reveal novel therapeutic strategies to treat NHO^17, 18^. Adult skeletal muscles contain two populations of stem/progenitor cells: *1)* satellite cells (SCs), residing within the myofiber under the myofiber basal lamina, which regenerate myoblasts and myocytes following injury and are as such the true muscle stem cells^22–24^, and *2)* fibro-adipogenic progenitors (FAPs) residing in the interstitial space between myofibers ^25^. Unlike SCs, FAPs are of mesenchymal origin and do not regenerate myoblasts^25^. Muscle repair following injury is a highly orchestrated process which involves the coordinated recruitment of both SCs and FAPs as well as macrophages. Upon muscle injury, FAPs proliferate transiently for the first 3 days after the injury in mice and then undergo apoptosis under the effect of tumor-necrosis factor (TNF) released by infiltrating C-C chemokine receptor-2 (CCR2)^+^ inflammatory monocyte / macrophages that also clear the apoptotic FAPs^26, 27^. Both FAPs and macrophages are essential to orchestrate and complete appropriate myogenic repair from SCs and the critical function of FAPs is thought to involve the secretion of appropriate growth factors and extracellular matrix which enable SC proliferation, myogenic differentiation and myofiber assembly^25, 28, 29^. As both muscle SCs and FAPs have osteogenic potential in vitro ^19, 21^, the question of what are the cells-of-origin of NHO is not resolved.

Lineage-tracing experiments using the Cre-loxP system have been undertaken to identify which progenitors contribute to the inheritable genetic form of heterotopic ossification called fibrodysplasia ossificans progressiva (FOP). Unlike NHO which are frequent after a severe CNS injury, FOP is an extremely rare disorder caused by a gain-of-function missense single nucleotide mutation in the bone morphogenetic protein (BMP) type I receptor gene *ACVR1*^30^, resulting in a switch in ligand preference with aberrant activation of BMP signaling by activin-A instead of BMPs^31^. Most lineage tracing experiments to identify the “cells-of-origin” of heterotopic ossifications were performed in mouse models of FOP either expressing a Cre recombinase-inducible *ACVR1*^R206H^ mutant that causes FOP in humans, expression of a BMP transgene, or the insertion of a BMP containing implant. In these mouse models of FOP, a variety of gene promoters have been used to drive Cre recombinase expression. Based on Cre recombinase driven by the *Pax7* (Paired Box 7) or *Cdh5* (Cadherin 5) gene promoters, which are specific of SCs and endothelial cells respectively, it was concluded that neither SCs nor endothelial cells were the cells-of-origin of heterotopic ossifications in FOP^32^. However most of the gene promoters used in similar studies (e.g. Glast/*Slc1a3*, *Scx*, *Mx1*, *Tie2,* CD133/*Prom1* gene promoters)^32–34^ were not exclusively expressed in mesenchymal progenitor cells and the conclusion that FOP heterotopic ossifications were derived from mesenchymal cells was inferred by elimination of other possible cell candidates. In mouse model of FOP driven by overexpression of a BMP4 transgene, it was recently shown that *Gli1* expressing cells were integrated in HO^34^. However, BMP4 is a very strong osteogenic protein and how artificially high BMP4 expression by means of a transgene is relevant to SCI-induced NHO remains to be demonstrated^17, 19^. Recently a Cre mouse driven by the *Pdgfra* gene promoter has been used to demonstrate that HO induced by BMP-2 supplemented Matrigel^35^ or by overexpressing FOP-causing *ACVR1*^R206H^ mutation^36^ were originated from mesenchymal cells expressing platelet-derived growth factor receptor-α (PDGFRα). However, while relevant models of FOP, artificial expression of osteogenic *ACVR1*^R206H^ or *ACVR1*^Q207D^ mutants or artificial overexpression of a BMP transgene or implantation of BMP-containing scaffolds utilized in these studies above are of little biological and clinical relevance to acquired NHO developing after severe CNS injuries because NHOs occur with high prevalence in genetically normal patients of a broad range of ethnicity. In a mouse model of traumatic HO induced by body burn and tenotomy, and a model of FOP induced by induced Cre recombinase activation of the *ACVR1*^Q207D^ activating mutation, it has been found that both burn-induced HO in the tendon and subcutaneous FOP-induced HO were derived from *Prrx1* gene expressing mesenchymal progenitors^37^. However, these HO models do not involve the severe CNS trauma which defines and triggers NHO^17, 19^. Furthermore, while FOP flare-ups develop in many different types of tissues associated with muscles (e.g. skeletal muscles, tendons, ligaments, fascia, and aponeuroses)^38^, possibly due to the dominant effect of activating mutations of *ACVR1*, NHO are mostly intramuscular in otherwise genetically normal patients^39^. Therefore, which progenitor cells within skeletal muscles are the “cells-of-origin” for NHO developing after a SCI remains to be established. To do so, we crossed a Cre-inducible fluorescent Zoanthus green (ZsGreen) reporter mouse strain with either the *Pax7-CreET2* enabling specific expression in SCs^40^, or with the *Prrx1-Cre* strain which specifically expresses Cre recombinase transgene in mesenchymal stem/progenitor cells under the control of the *Prrx1* gene enhancer^41^. Finally, we sorted CD56^+^ myogenic progenitors and PDGFRα-expressing mesenchymal progenitors from human skeletal muscle surrounding NHO after surgical excision and studied their osteogenic differentiation capacity *in vitro* and *in vivo* in an ectopic bone model in immunodeficient mice.

## RESULTS

### ZsGreen^+^ cells identify SCs and FAPs in the muscles of *Pax7*-CreERT2 and *Prrx1*-Cre mice respectively

To determine whether NHO is derived from muscle SCs and FAPs, we first assessed the specificity and efficiency of ZsGreen reporter expression in *Pax7*-CreERT2; *Rosa26*-LoxP-STOP-LoxP-ZsGreen mice (abbreviated as *Pax7*^ZsG^ thereafter) and *Prrx1*-Cre; *Rosa26*-LoxP-STOP-LoxP-ZsGreen mice (abbreviated as *Prrx1*^ZsG^) (Fig. 1a). CreERT2 was activated by daily gavage with tamoxifen for 4 days after which mice were left to rest for an additional 2 weeks. The right hind limb hamstring muscle was then injected with cardiotoxin (CDTX) to cause muscle injury and initiate muscle regeneration. Fourteen days post-injury, muscles were harvested and dissociated for analyses of single muscle cell suspension by flow cytometry (Fig. 1b). Within the CD45^−^ and hematopoietic lineage (Lin: Ter119, CD3ε, CD45R, CD11b, Gr1)-negative non-hematopoietic cell gate, endothelial cells were identified as CD31^+^ and stem cell antigen-1 (Sca1)-positive, SCs identified as CD31^−^ CD34^+^ Sca1^−^ integrin α7 (ITGA7)^+^ PDGFRα^−^ and FAPs identified as CD31^−^ CD34^+^ Sca1^+^ ITGA7^−^ PDGFRα^+^ (Fig. S1) in accordance with previous literature^23, 25, 42^. The frequencies of SCs (CD31^−^ CD34^+^ Sca1^−^ ITGA7^+^ PDGFRα^−^), FAPs (CD31^−^ CD34^+^ Sca1^+^ ITGA7^−^ PDGFRα^+^) and endothelial cells (CD31^+^ Sca1^+^) were similar between the CDTX-injured and non-injured muscles in both *Pax7*^ZsG^ and *Prrx1*^ZsG^ strains 14 days post-injury (Table S1).

**Fig. 1.**
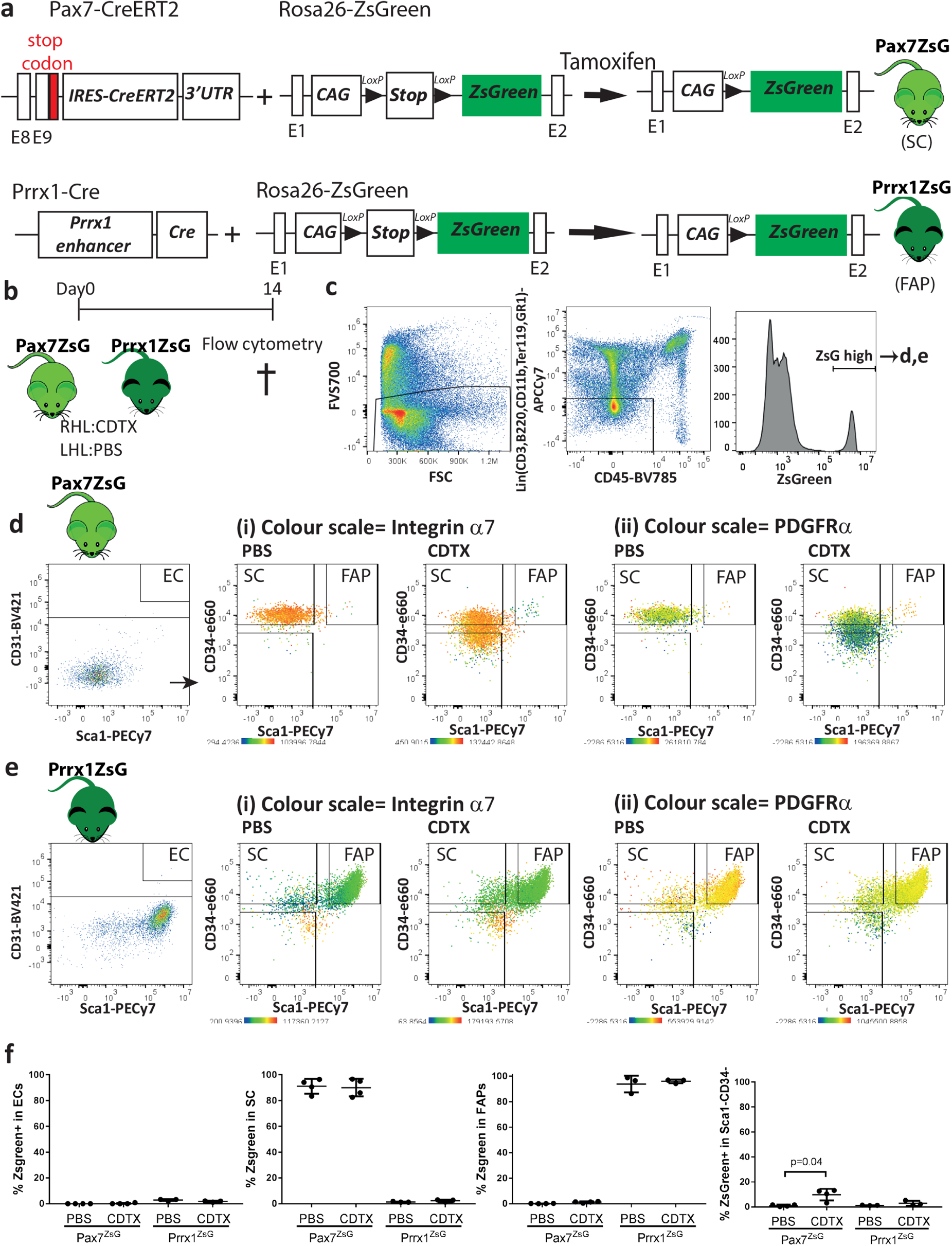
ZsGreen labels SCs and FAPs in skeletal muscles of Pax7^ZsG^ and Prrx1^ZsG^ mice respectively. **a** Schematic representation of the tamoxifen-inducible Cre-dependent ZsGreen reporter induction. **b** *Pax7*^ZsG^ (n=4/group) and *Prrx*1^ZsG^ (n=3/group) mice were injected with CDTX in the hamstring muscle of right hind limb (RHL) and PBS in the hamstring muscle of left hind limb (LHL). Muscle cells were isolated 14 days after injection and subsequently stained for surface cell markers. **c** After gating on forward/side scatters and FVS700 dead cell exclusion, ZsGreen-high cells were gated from Lin^−^ CD45^−^ non-hematopoietic population in both **d** *Pax7*^ZsG^ and **e** *Prrx1*^ZsG^ mice and further sub-grouped into CD31^+^Sca1^+^(EC), CD31^−^Sca1^−^CD34^+^(SCs), CD31^−^Sca1^+^CD34^+^ (FAPs) and Sca1^−^ CD34^−^ populations. The color indicates the expression level of (**i**) integrin α7 and (**ii**) PDGFRα. Yellow-orange: high expression. Green: low expression. **f** Frequencies of ZsGreen^+^ cells in Sca1^+^CD31^+^ ECs, CD31^−^ Sca1^−^ CD34^+^ ITGA7^+^ PDGFRα^−^ SCs and CD31^−^ Sca1^+^ CD34^+^ ITGA7^−^ PDGFRα^+^ FAPs populations from muscles of *Pax7*^ZsG^ and *Prrx1*^ZsG^ nice following the gating strategy in Figure S1. Each dot represents a separate mouse and bars are mean ± SD. Significance was analyzed by one-way ANOVA with Tukey’s multiple comparison test.

We subsequently gated the ZsGreen^+^ cells within the hamstring muscles of these mice (Fig. 1c). In the vehicle-injected muscles of *Pax*7^ZsG^ mice, all ZsGreen^+^ cells had the typical SC phenotype (CD45^−^ Lin^−^ CD31^−^ CD34^+^ Sca1^−^ ITGA7^+^ PDGFRα^−^) with no ZsGreen^+^ cell in the endothelial gate and very few in the FAP gate (Fig. 1d - PBS). In the CDTX-injured muscle, the distribution of ZsGreen^+^ cells was similar but with the emergence of CD45^−^ Lin^−^ CD34^−^ Sca1^−^ ITGA7^+^ PDGFRα^−^ cells derived from ZsGreen^+^ SCs (Fig. 1d - CDTX). These cells are likely differentiating myoblasts.

Conversely, in the vehicle-injected contralateral muscle of *Prrx1*^ZsG^ mice, most ZsGreen^+^ cells had the typical FAP phenotype with no ZsGreen^+^ cell in the endothelial gate and very few in the typical SC gate (Fig. 1e PBS). The phenotypes of ZsGreen^+^ cells were the same in the CDTX-injured muscle of *Prrx1*^ZsG^ mice (Fig. 1e - CDTX).

In addition to the high specificity of these lineage tracing strategies, the recombination efficiency was also very high with over 90% of SCs and FAPs labelled in the muscle of *Pax7*^ZsG^ and *Prrx1*^ZsG^ strains respectively (Fig. 1f). Importantly, Cre-mediated recombination in *Pax7*^ZsG^ and *Prrx1*^ZsG^ mice did not affect subsequent NHO development compared to C57BL/6 mice as measured by micro-computerized tomography (µCT) or by hematoxylin-eosin staining, which clearly showed the presence of necrotic and fibrotic muscle together with inflammatory infiltrate and sporadic bony nodules 28 days after injury in both strains as we previously reported in C57BL/6 mice^13, 20, 21, 43^ (Fig. S2). Altogether, these experiments confirmed that ZsGreen was specifically expressed in the targeted muscle cell populations in both *Pax7*^ZsG^ and *Prrx1*^ZsG^ strains without altering NHO development thus illustrating their suitability for lineage tracing of SCs and FAPs in vivo.

**Fig. 2.**
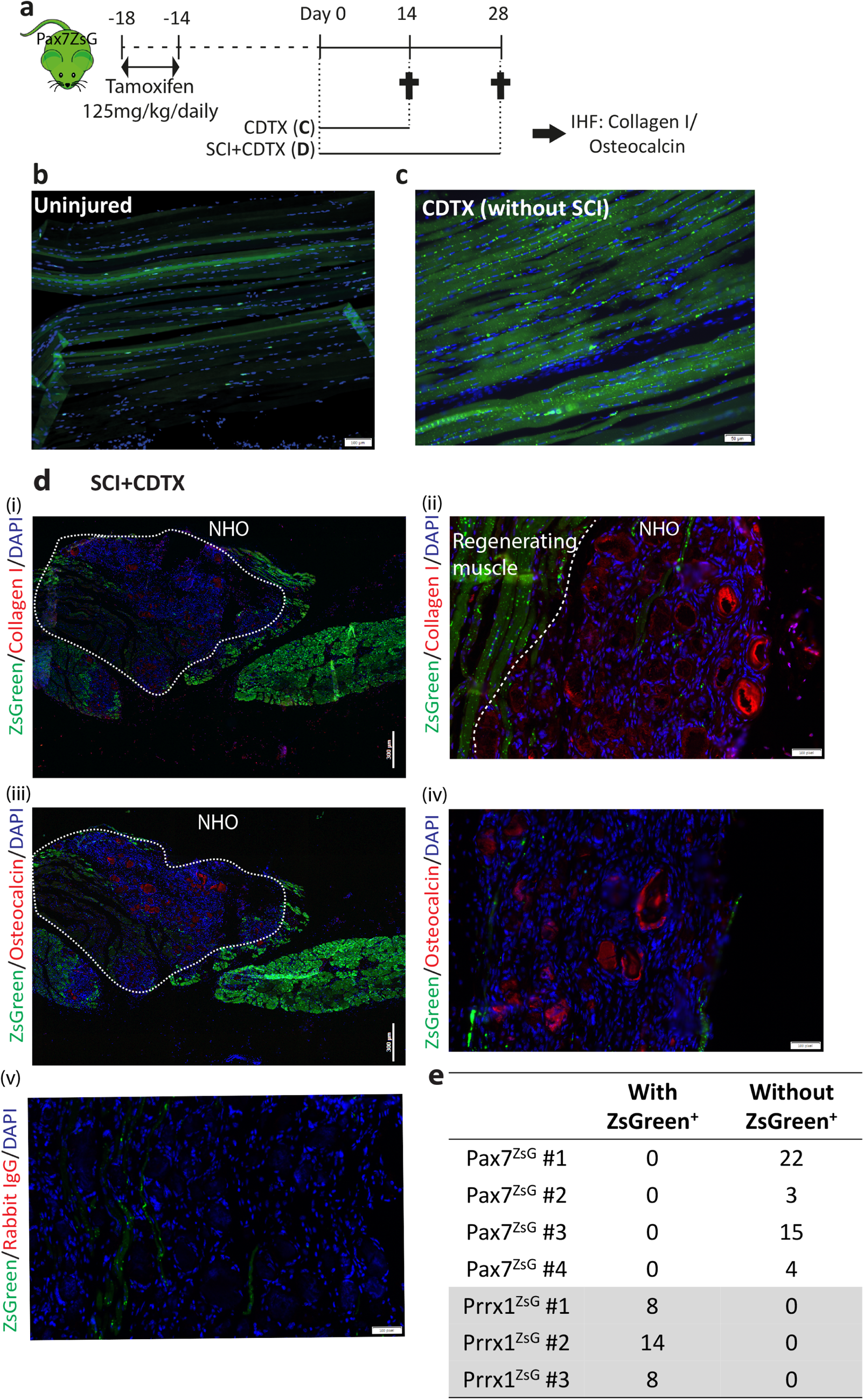
NHO are not derived from *Pax7* expressing SCs. **a** ZsGreen expression in *Pax7*^ZsG^ mice was induced by tamoxifen treatment for 4 days. Two weeks after tamoxifen treatment, mice received an intramuscular injection of CDTX with or without SCI. Muscle samples were harvested 14 and 28 days post injury respectively. Representative IHF images illustrating the distribution of ZsGreen^+^ SC-derived cells in **b** uninjured muscle and **c** regenerated injured muscle 14 days post CDTX injection in *Pax7*^ZsG^ mice without SCI (n=3 mice/group). Scales bars: **b** 100μm, **c** 50μm. **d** Representative images from *Pax7*^ZsG^ mice with SCI and CDTX-mediated muscle injury 28 days post-surgery. (i) IHF staining illustrating that ZsGreen^+^ cells are present amongst areas of regenerating muscle and largely absent from areas of NHO development. White dashed lines indicate the boundary between regenerating muscle and fibrotic area containing NHO stained red (i-ii) collagen I and (iii-iv) osteocalcin. Nuclei were stained by DAPI staining (blue). Negative control was performed using rabbit isotype IgG (iv). Scale bars: **d (i, iii)** 300 μm, **(ii, iv, v)** 50μm. **e** Number of osteocalcin^+^ NHO intercalated with ZsGreen^+^ cells or without ZsGreen^+^ cells in both *Pax7*^ZsG^ (n=4 mice, total 44 osteocalcin^+^ NHO counted) and *Prrx*1^ZsG^ mice (n=3 mice, total 30 osteocalcin^+^ NHO counted). Statistical difference was calculated using a Fisher Exact test.

### NHOs are not derived from *Pax7* expressing SCs

ZsGreen expression was then examined on frozen longitudinal sections of un-injured muscle, repaired muscle and NHO from *Pax7*^ZsG^ mice (Fig. 2a). In non-injured muscles from *Pax7*^ZsG^ mice, ZsGreen was expressed in a discrete population of small cells distributed along myofibers typical of SCs (Fig. 2b). Some myofibers were also ZsGreen^+^ likely reflecting their natural turn-over from ZsGreen^+^ SCs over a period of 4 weeks after tamoxifen induction. Fourteen days after CDTX-induced muscle injury, in the absence of SCI, all myofibers were ZsGreen^+^ as anticipated, illustrating muscle regeneration from ZsGreen^+^ SCs (Fig. 2c).

In the cohort of mice that underwent spinal cord transection (SCI) and CDTX-mediated muscle injury, frozen sections were stained with specific anti-collagen I (Fig. 2d(i-ii)) and anti-osteocalcin antibodies (Fig. 2d(iii-iv)). Example of non-immune IgG negative control for anti-collagen I and anti-osteocalcin antibodies is shown in Fig. 2d(v). Immuno-histofluorescence (IHF) confirmed ZsGreen^+^ cells within areas of regenerating muscle which contained neo-formed ZsGreen^+^ myofibers. Most importantly, ZsGreen^+^ cells were largely absent amongst areas of fibrotic muscle and NHO nodules. ZsGreen^+^ cells were not intercalated among the collagen I^+^ bone matrix or osteocalcin^+^ osteoblasts on NHO nodules. Quantification of NHO through four different *Pax7*^ZsG^ mice showed none of the 44 osteocalcin^+^ NHO contained ZsGreen^+^ cells (Fig. 2e). This demonstrates that NHO following SCI are not derived from ZsGreen^+^ SCs.

### NHOs are derived from *Prrx1* expressing FAPs

ZsGreen expression was also examined on frozen longitudinal sections of un-injured muscle, repaired muscle and NHO from *Prrx1*^ZsG^ mice (Fig. 3a). In un-injured muscle, reticulated ZsGreen^+^ cells were scattered in the interstitium along myofibers (Fig. 3b), a typical distribution and morphology of FAPs^25^. Importantly, in the regenerating CDTX-injured muscle (without SCI) ZsGreen^+^ cells were distributed similarly and most importantly, they did not contribute to neoformation of myofibers (Fig. 3c) in concordance with the literature^25^.

**Fig. 3.**
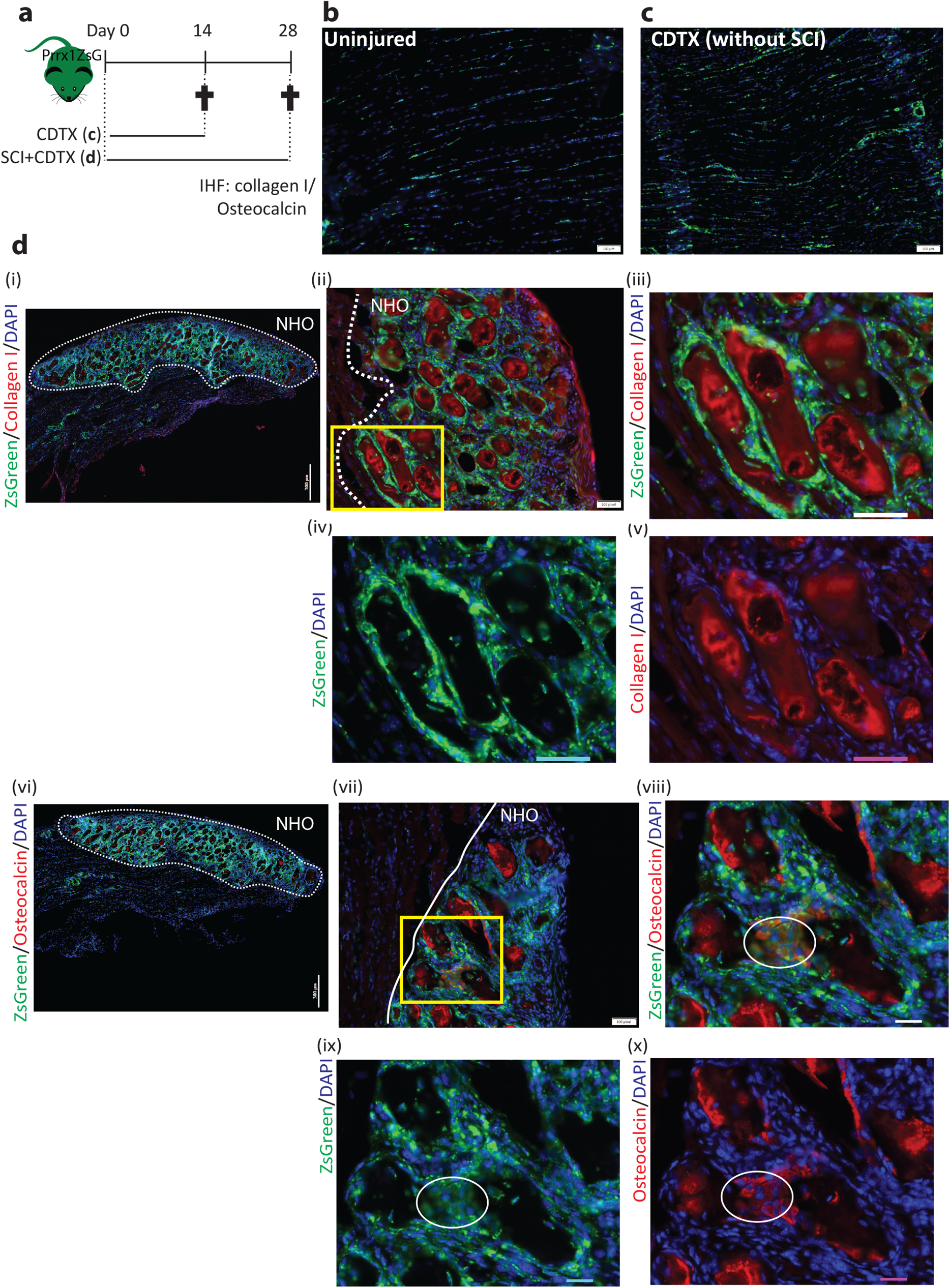
NHO are derived from *Prrx1* expressing FAPs. **a** *Prrx*1^ZsG^ mice received an intramuscular injection of CDTX with or without SCI, and muscle samples were harvested at indicated time points and processed for IHF. Representative images illustrating the distribution of ZsGreen^+^ FAP-derived cells in **b** un-injured muscle and **c** regenerated injured muscle 14 days post CDTX injection in *Prrx1*^ZsG^ mice without SCI (n=3 mice/group). **d** Representative images from *Prrx1*^ZsG^ mice with SCI and CDTX-mediated muscle injury 28 days post-surgery: IHF illustrating the co-localization of ZsGreen^+^ FAP-derived cells with i-v) collagen I^+^ matrix (red) or (**vi-x**) osteocalcin^+^ osteoblasts (red). (**iii-v**) are enlarged images of yellow box in (**ii**) whereas (**viii-x**) are enlarged image of yellow box in (**vii**). White dashed line indicates the boundary between regenerating muscle and fibrotic area containing NHO. Nuclei stained in DAPI (blue). White circles indicate osteocalcin^+^ osteoblasts which also express ZsGreen. Scale bars: **b-c** 100μm; d (i) 300 μm (**ii, vii**) 100 μm (**iii-v, viii-x**) 50μm.

The *Prrx1*^ZsG^ mice that underwent SCI and CDTX-mediated muscle injury also developed NHO (similar to *Pax7*^ZsG^ mice that underwent the same treatment (Fig.S2). However, in sharp contrast to *Pax7*^ZsG^ mice, all fibrotic areas in the SCI+CDTX-injured muscles of *Prrx1*^ZsG^ mice were intensely labeled by ZsGreen, particularly around collagen I^+^ and osteocalcin^+^ NHO nodules (Fig. 3d(ii-x)). We counted 30 osteocalcin^+^ NHO from three *Prrx1*^ZsG^ mice and all of them were intercalated with ZsGreen^+^ cells (Fig. 2e). At higher magnification, differentiating osteocalcin^+^ osteoblasts (red) were also ZsGreen^+^ (Fig. 3d(vii-x) circles), clearly demonstrating that NHO are derived from *Prrx1*-expressing mesenchymal progenitor cells rather than SCs.

### Mesenchymal *Prrx1*^+^ cells do not circulate in blood after SCI

Parabiosis models have been previously used to track circulating cells that contribute to heterotopic ossification (HO). An ubiquitous green fluorescent protein (GFP) reporter mouse was joined with a wildtype mouse, with the wildtype mouse receiving an Achilles tendon tenectomy and dorsal burn injury to induce HO formation. Circulating GFP^+^ cells from the GFP-expressing parabiont contributed to HO developing in the wild-type parabiont 28 days after tenectomy and burn injury^44^.

However, a more recent parabiosis study suggests otherwise: HO induced by inserting BMP2-supplemented Matrigel matrix did not contain cells derived from PDGFRα^+^ FOPs recruited from the other parabiont via the shared circulation suggesting that the cells-of-origin of BMP2-induced HO are not recruited via the circulation^35^.

In order to clarify the dichotomy and these two previous publications, we first investigated whether SCI activates expression of osteogenic BMPs and BMP signaling in injured muscles. SCI did not increase the expression of *Bmp2*, *Bmp4* or *Bmp7* RNA in paraplegic hindlimb muscles (Fig.S3a). On the contrary, either SCI or muscle injuries decreased the expression of the osteogenic BMPs. We also investigated the effect of a daily treatment with LDN-193189, a potent inhibitor of BMP type I receptor serine kinases which has been shown to be effective at inhibiting HO development in the mouse FOP model driven by the *ACVR1*^Q207D^ missense mutation^45^ and in the burn-induced rat model of ossifying tendinopathy^46^. While LDN-193189 significantly inhibited mineralization of mouse bone marrow-derived mesenchymal stromal cells (BM-MSCs) cultured in osteogenic conditions (Fig. S3b), mouse treatment with LDN-193189 either in the first two weeks of the injury or during the maturation phase of NHO between weeks 2 and 3 post-injury, had no effect on NHO development (Fig. S3c-d). As BMPs are known to induce endochondral ossification, we stained sections of CDTX-injured muscles at days 7,14 and 21 following SCI with safranin O to span the time course of NHO development in mice (Fig. S4). Although occasional mast cells brightly stained by safranin O could be detected in the developing NHO, there was no evidence of cartilage matrix in any NHO examined at any times-point (n = mice per time-point). Therefore, unlike the *ACVR1*^Q207D^ FOP model and the burn-induced ossifying tendinopathy model, the BMP signaling inhibitor LDN-193189 was unable to reduce NHO development after SCI suggesting NHO development following SCI is not as BMP-dependent as the two other processes of HO development and does not involve endochondral ossification.

We next investigated whether *Prrx1*^+^ mesenchymal cells at the origin of NHO could be derived from the circulation. As parabiosis experiments are forbidden for ethical reasons in Australia, we investigated whether ZsGreen^+^ cells were detectable in the circulation of *Prrx1*^ZsG^ mice 1, 2, 3 and 7 days following SCI and CDTX-mediated muscle injury (Fig. 4b-c). Flow cytometry on blood nucleated cells showed that there was a ZsGreen^low^ population that mostly included CD45^+^ CD11b^+^ F4/80^+^ monocytes and CD45^+^ CD11b^+^ F4/80^−^ populations (Fig. 4c) but the intensity of ZsGreen fluorescence in these phagocytes was two full log10 units lower than the ZsGreen fluorescence intensity of FAPs in the muscle (Fig. 4d). Therefore, it is likely that the low ZsGreen fluorescence of blood monocytes and granulocytes is due to phagocytosis of ZsGreen (possibly packaged in extracellular vesicles^47, 48^) produced by *Prrx1*-expressing mesenchymal progenitor cells and their progenies. Importantly, despite analyzing over 10^6^ blood leukocytes per mouse, we could not detect any circulating CD45^−^ Lin^−^ CD31^−^ Sca1^+^ mesenchymal progenitor cells or cells with ZsGreen fluorescence intensity as high as FAPs in the muscle of *Prrx1*^ZsG^ mice (Fig. 4b,d). As another potential source of “mobilized” mesenchymal cells is the bone marrow, we transplanted 90,000 stromal cells enriched from the bone marrow of *Prrx1*^ZsG^ mice by magnetic activated cell depletion of CD45^+^ leukocytes and Ter119^+^ erythroid cells using the EasySep mouse stromal cell enrichment kit (Stem Cell Technologies). Following this enrichment step, 11.7% of these bone marrow stromal cells were ZsG^bright^ with the classic CD45^−^ Lin^−^ CD31^−^ Sca1^+^ mesenchymal progenitor cell phenotype. These cells were transplanted i.v. into three C57BL/6 recipients 24 hours post-SCI and CDTX muscle injury of the recipients. Muscles were analyzed by histofluorescence 21 days later and none of the CDTX-injured muscles contained any ZsGreen-positive cell (result not shown). Therefore, while we cannot completely exclude the possibility that some of the *Prrx1*-expressing ZsGreen^+^ cells associated with NHO following SCI may come from the circulation, these would be extremely rare compared to the abundance of *Prrx1*-expressing ZsGreen^+^ FAPs already present in the muscle prior to injury. Therefore, NHO are most likely derived from muscle-resident FAPs.

**Fig. 4.**
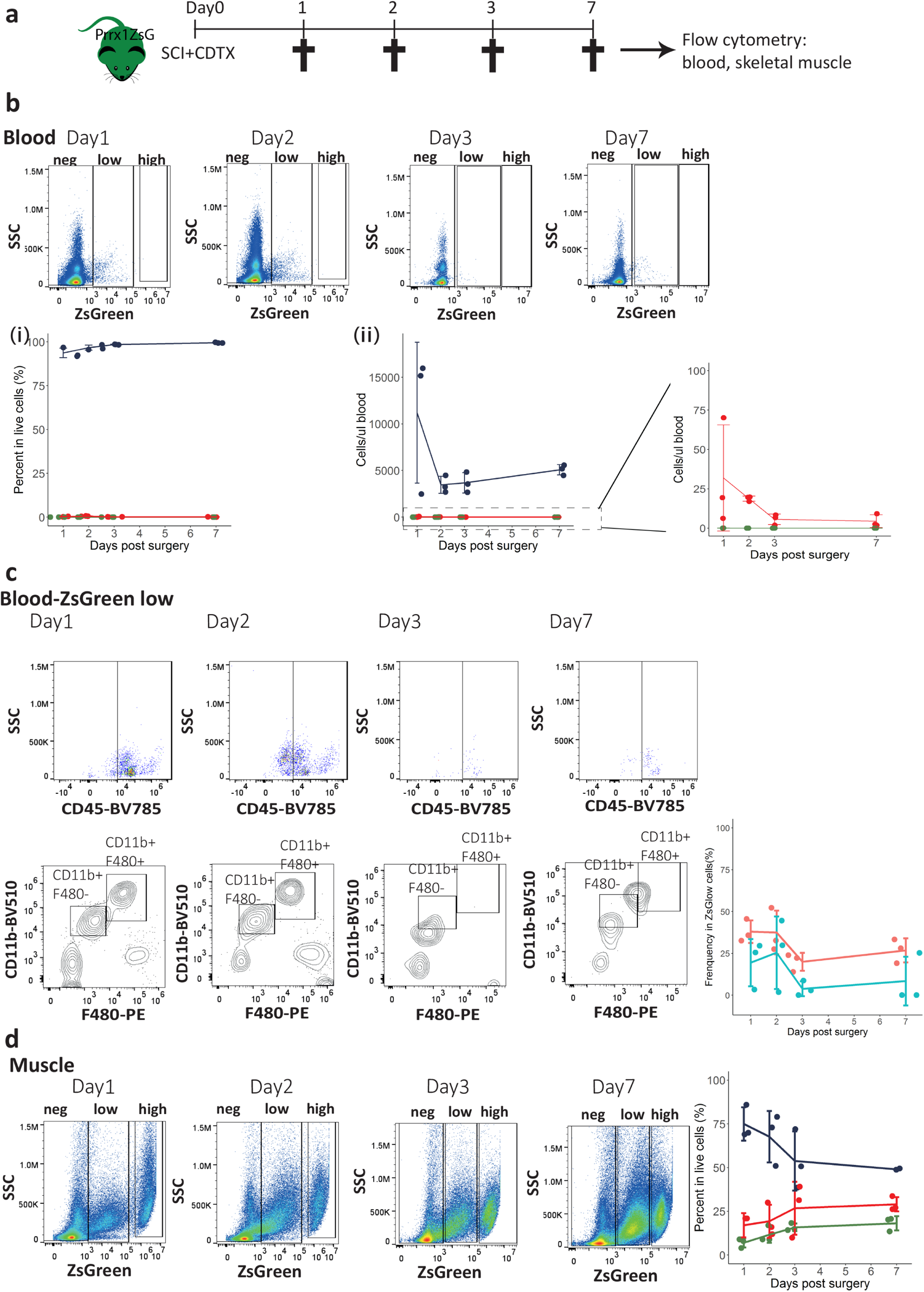
Absence of ZsGreen^high^ mesenchymal cells in the circulation of *Prrx1*^ZsG^ mice after SCI and muscle injury. **a** *Prrx*1^ZsG^ mice received SCI and an intramuscular injection of CDTX. Peripheral blood and skeletal muscles were collected at 1, 2, 3 and 7 days post-surgery and analyzed by flow cytometry (n=3 mice/group). **b** FVS700-Live cells from peripheral blood were gated based on the intensity of ZsGreen fluorescence into ZsGreen negative (blue lines), low (red lines) and high (green lines). Frequency of these three populations were represented as (**i**) frequency of live cells in blood and (**ii**) numbers of cells per μl blood. **c** The ZsGreen-low cells were further gated for expression of CD11b and F480. The frequency of CD11b^+^ F4/80^+^ monocytes (red line) and CD11b^+^ F4/80^−^ granulocytes (blue line) among circulating ZsG^low^ cells was plotted. **d** Live cells from injured muscle from the same *Prrx1*^ZsG^ mice were used as reference for ZsGreen fluorescence intensity, confirming the presence of numerous ZsGreen^high^ cells in muscle. ZsGreen negative (blue lines), low (red lines) and high (green lines). Each dot represents a separate mouse. Bars represent means ± SD. There was no significant difference between the time points as determined by one-way ANOVA with Tukey’s multiple comparison test.

### Spinal cord injury causes PDGFRα up-regulation on FAPs irrespective of muscle injury

To better understand the effect of the SCI on muscle FAPs, we followed PDGFRα expression on muscle cells. Surprisingly, PDGFRα was significantly upregulated on CD45^−^ Lin^−^ CD31^−^ Sca1^+^ CD34^+^ FAPs at 7 and 14 days after SCI both in CDTX-injured and contralateral non-injured muscles (Fig. S5a-c). However, no expression of PDGFRα was noted on CD45^−^ Lin^−^ CD31^−^ Sca1^−^ CD34^+^ SCs in any of the experimental conditions tested (Fig. S5d). Although it remains to be determined whether this up-regulation of PDGFRα has a functional role in driving NHO development, this finding suggests that SCI is able to alter or reprogram FAP function as detected by PDGFRα expression.

### Spinal cord injury leads to reduced apoptosis and persistent proliferation of FAPs following muscle injury

Sequential induction of FAP proliferation followed by their apoptosis is crucial for effective muscle repair as either increased proliferation or decreased apoptosis can result in over-expansion of FAPs in the regenerating muscle resulting in fibrosis^27^. To examine whether the dynamics of FAP proliferation and apoptosis are perturbed during NHO formation, we first compared by flow cytometry the frequencies of apoptotic cells within FAP and SC populations three days post injuries, a time point at which FAP apoptosis peaks in the regenerating muscle^27^ (Fig. 5a). We stained muscle cell suspension with annexin V (AnnV) to detect early stage of apoptosis together non-membrane permeable DNA dye with 7-amino-actinomycin D (7AAD) to identify live cells (AnnV^−^ 7AAD^−^), apoptotic cells (AnnV^+^ 7AAD^−^) and post-apoptotic dead cells (AnnV^+^ 7AAD^+^). Within the Lin^−^ CD45^−^ CD31^−^ Sca1^+^ CD34^+^ ITGA7^−^ FAP population, muscle injury alone (Sham+CDTX) led to 17% reduction in live FAPs and 3.2-fold increase of apoptotic FAP frequency compared to naïve muscle as previously reported^27^ (Fig. 5a(i)). However, in mice that had undergone SCI together with muscle injury, the frequencies of live and apoptotic FAPs was reversed to those found in naive mice. In respect to the Lin^−^ CD45^−^ CD31^−^ Sca1^−^ CD34^+^ ITGA7^+^ SC population, CDTX injury led to an 81% reduction of live SCs and 2.6-2.7 fold increase in dead SCs regardless of presence or absence of SCI (Fig. 5a(ii)).

**Fig. 5.**
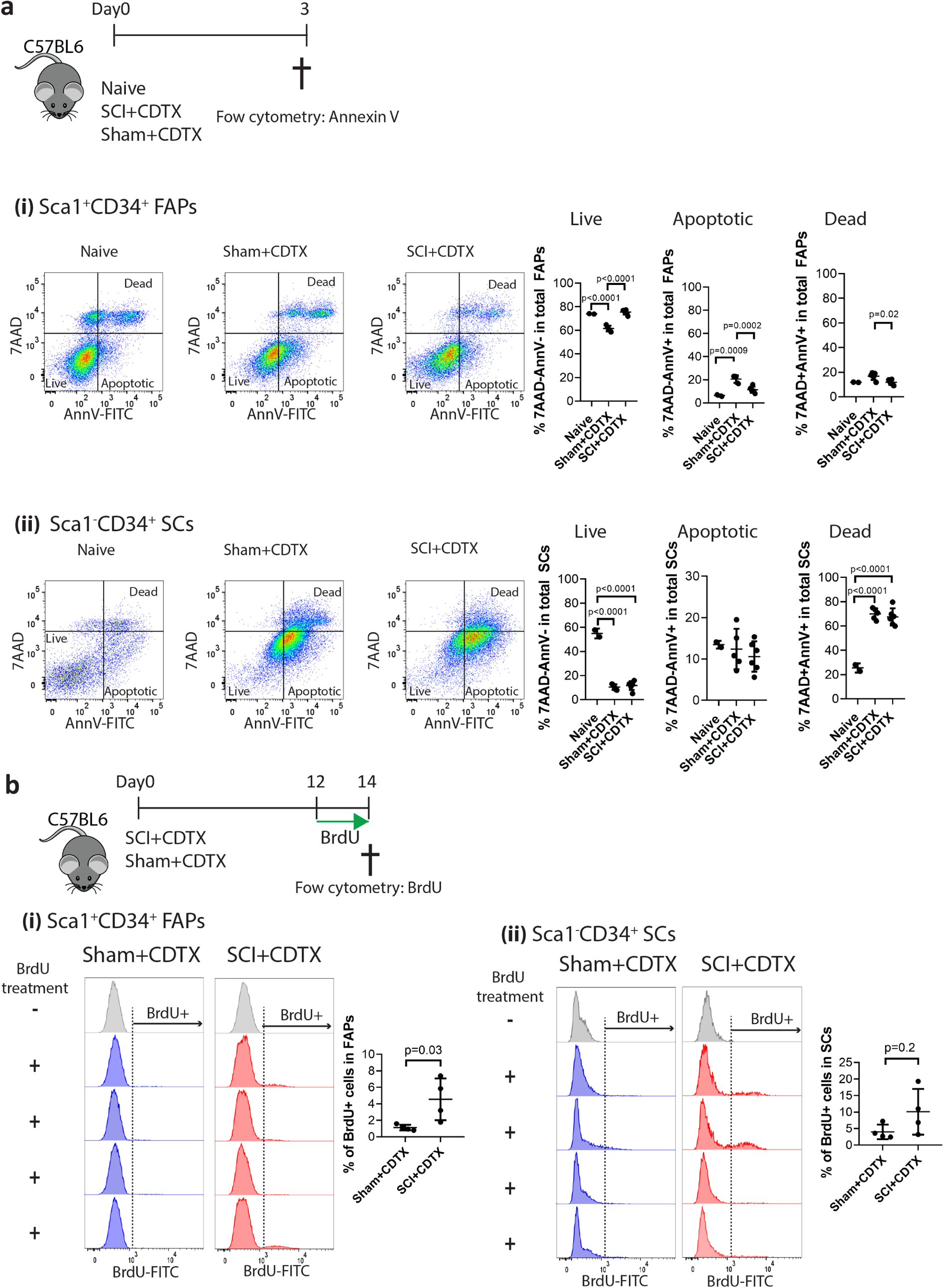
SCI leads to decreased apoptosis and persistent proliferation of FAPs in injured muscles. **a** Naïve C57BL/6 mice underwent SCI or Sham surgery plus intramuscular injection of CDTX. Muscle cells were isolated 3 days later. Apoptotic cells were subsequently analyzed by annexin V (AnnV) and 7-amino-actinomycin D (7AAD) staining by flow cytometer. (i) CD45 Lin^−^ CD31^−^Sca1^+^ CD34^+^ ITGA7^−^ FAPs and (ii) CD45 Lin^−^ CD31^−^Sca1^−^ CD34^+^ ITGA7^+^ SCs were gated. Ann V and 7AAD staining further distinguished cells as live (7AAD^−^AnnV^−^), apoptotic (7AAD^−^AnnV^+^) and post-apoptotic dead (7AAD^+^AnnV^+^) in both FAP and SC populations. Percentage of live, apoptotic and dead cells in total FAP or SC populations are presented as mean ± SD (n=2,5,6 in naïve, Sham+CDTX, SCI+CDTX respectively). Each dot represents a separate mouse. Significance was calculated by one-way ANOVA with Tukey’s multiple comparison test. **b** C57BL/6 mice underwent SCI or Sham surgery plus intramuscular injection of CDTX. Mice were supplemented with drinking water containing BrdU together with BrdU i.p. injection (twice daily) from day 12-14. One SCI+CDTX and 1 Sham+CDTX mouse were not treated with BrdU and used as negative control for anti-BrdU staining. On day14, muscle cells were isolated and BrdU staining was analyzed in CD45 Lin^−^ CD31^−^Sca1^+^ CD34^+^ ITGA7^−^ FAPs and CD45 Lin^−^ CD31^−^Sca1^−^ CD34^+^ ITGA7^+^ SCs by flow cytometry. Percentage of BrdU^+^ cells in total FAP or SC populations are presented as mean ± SD. Each dot represents a separate mouse. Significance was calculated by two-sided Mann-Whitney test.

We next measured muscle progenitor proliferation by in vivo 5-bromo-2’-deoxyuridine (BrdU) incorporation 14 days post surgeries. To ensure sufficient incorporation of BrdU into newly synthesized DNA during cell proliferation, mice were treated with BrdU two days prior to tissue harvest. The percentage of proliferative BrdU^+^ FAPs was 4.08-fold higher in the SCI+CDTX group compared to the Sham+CDTX group (Fig. 5b(i)) while there was no significant difference in the proliferation of SCs (Fig. 5b(ii)). These results confirm that SCI deregulates the coordinated proliferation and apoptosis of FAPs in injured muscles.

### Human PDGFRα^+^ cells from muscles surrounding NHO support both in vitro and in vivo bone formation

To validate the involvement of muscle-resident mesenchymal cells in the human NHO pathology, we collected surgical residues of NHO after resection surgery in twelve SCI, TBI and stroke patients. Cells were isolated from the muscle tissue surrounding the resected NHOs using mechanical dissociation and enzymatic digestion and expanded in culture. PDGFRα^+^ and CD56^+^ cells were then isolated by fluorescence activated cell sorting (Fig. 6a). PDGFRα^+^ population displayed a classical mesenchymal phenotype CD31^−^ CD45^−^ CD73^+^ CD90^+^ CD105^+^ (Fig. 6b(i)). Interestingly PDGFRα^+^ cells showed a heterogeneous expression of CD34, a common marker of hematopoietic and angiogenic progenitor cells (from 2% to 46.1%; Fig. 6b(ii)). As previously described ^49^, CD56^+^ cells were CD73^+^ CD90^+^ CD105^+^ (Fig. 6b(ii)) and expressed the myogenic regulatory transcription factors *MYF5* and *MYOD1* (Fig. 6b(iii)) whereas sorted PDGRα^+^ cells did not.

**Fig. 6.**
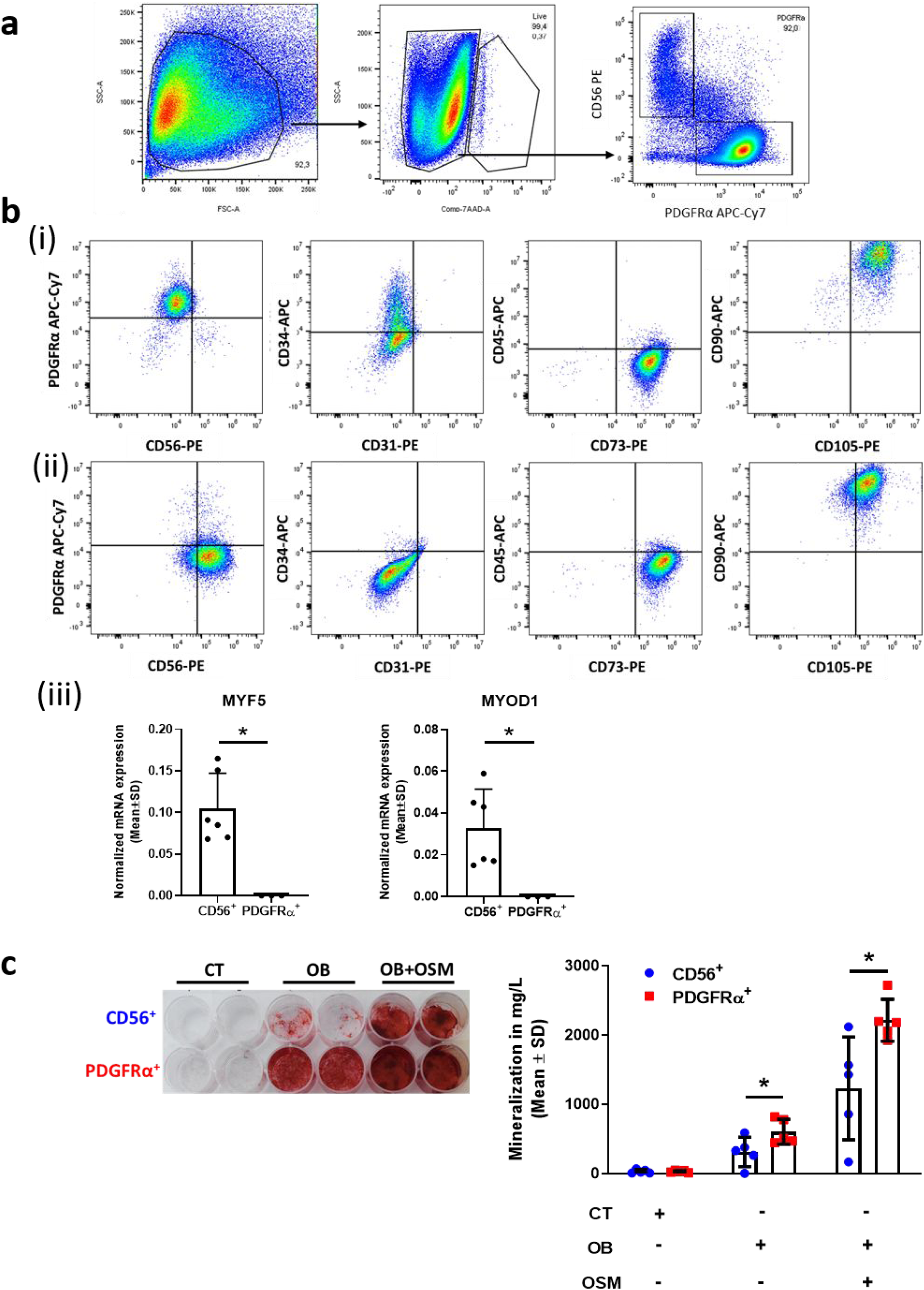
Human PDGFRα^+^ cells isolated from muscles surrounding NHO support *in vitro* bone formation. **a** Flow cytometry gating strategy of PDGFRα^+^ and CD56^+^ cell subpopulations isolated from muscle surrounding NHO. **b** Representative surface markers characterization by flow cytometry: CD56, PDGFRα, CD31, CD34, CD45, CD73, CD90, and CD105, (**i**) CD56^+^ population, (**ii**) PDGFRα^+^ population, (**iii**) Normalized mRNA expression of MYF5 and MYOD1 by qRT-PCR expressed as mean ± SD (CD56^+^ n=6; PDGFRα^+^ n=3). **c** *In vitro* osteoblastic differentiation assay seeded with CD56^+^ or PDGFRα^+^ cells isolated from muscles surrounding human NHO. **(i**) All cells were cultured in control medium (CT) or osteogenic medium alone (OB) or were supplemented with human OSM (100 ng/ml) (OB + OSM) for 14 days followed by Alizarin Red S staining. **c**(**ii**) Quantification of calcium mineralization expressed as mean ± SD (n=5). *p<0.05, two-sided non-parametric Mann Whitney U test.

*In vitro* osteogenic differentiation assays showed that PDGFRα^+^ cells exhibited higher osteoblastic differentiation capacity compared to CD56^+^ cells (Fig. 6c) and osteogenic differentiation was enhanced by the addition of 100 ng/ml recombinant human OSM (Fig. 6c), a key inflammatory mediator of NHO formation, as we previously reported ^21^.

We then investigated whether cells sorted from muscles surrounding human NHO were able to support *in vivo* heterotopic bone formation in immunodeficient mice. PDGFRα^+^ and CD56^+^ cells were independently seeded into plasma clotted hydroxyapatite/calcium phosphate scaffolds and implanted subcutaneously into the back of nude mice (Fig. 7a). Unseeded plasma clotted scaffolds were used as negative control and BM-derived mesenchymal stromal cells (BM-MSCs) seeded plasma clotted scaffolds as positive controls. Fifteen weeks after implantation, scaffolds were collected for histological analysis. As expected, no bone tissue was observed in the control plasma group (Fig. S6a(i)) while all BM-MSC seeded implants exhibited bone matrix and hematopoietic foci as detected by hematoxylin, eosin and safran staining (HES) (Fig. S6a(ii)). Six out of 11 implants (54.5%) with NHO muscle PDGFRα^+^ cells showed mature bone matrix containing osteocytes together with a hematopoietic marrow demonstrating the formation of an ectopic bone with functional hematopoietic BM (Fig. 7b and 7c(i)) as evidenced by the presence of numerous mature megakaryocytes (Fig. S6b). In contrast, only 12.5% of scaffolds seeded with CD56^+^ cells contained bone matrix deposition alone and 12.5% showed both bone matrix development and hematopoietic colonization (Fig. 7b and 7c(i)) suggesting a much higher osteogenic potential of PDGFRα^+^ mesenchymal cells derived from human muscle surrounding NHO. In order to analyze the origin of bone forming cells in these implants, the expression of human-specific lamin A/C expression was investigated by immunohistochemistry. Fig. 7c(ii) highlights that very few human cells were observed within CD56^+^ cell seeded scaffolds, whereas PDGFRα^+^ cell seeded implants exhibited numerous human lamin A/C^+^ osteocytes within the bone matrix (Fig. 7c(ii)), demonstrating that human PDGFRα^+^ cells actively participate in the formation of heterotopic bone. Of note, no human cells were detected within hematopoietic foci, confirming that hematopoietic cells that colonized these heterotopic bone formations were from murine origin (Fig. S6b). Finally, PDGFRα^+^ cell implants exhibited numerous osterix^+^ osteoblasts; most of them were located nearby hydroxyapatite particles and within the bone matrix (Fig. 7d). Thus, mesenchymal PDGFRα^+^ cells isolated from muscles surrounding human NHOs have a substantially higher capacity to support mature hemogenic bone formation compared to myogenic CD56^+^ cells. These data are concordant with the hypothesis that muscle-resident PDGFRα^+^ cells are key actors in the onset of NHO in both humans and mice.

**Fig. 7.**
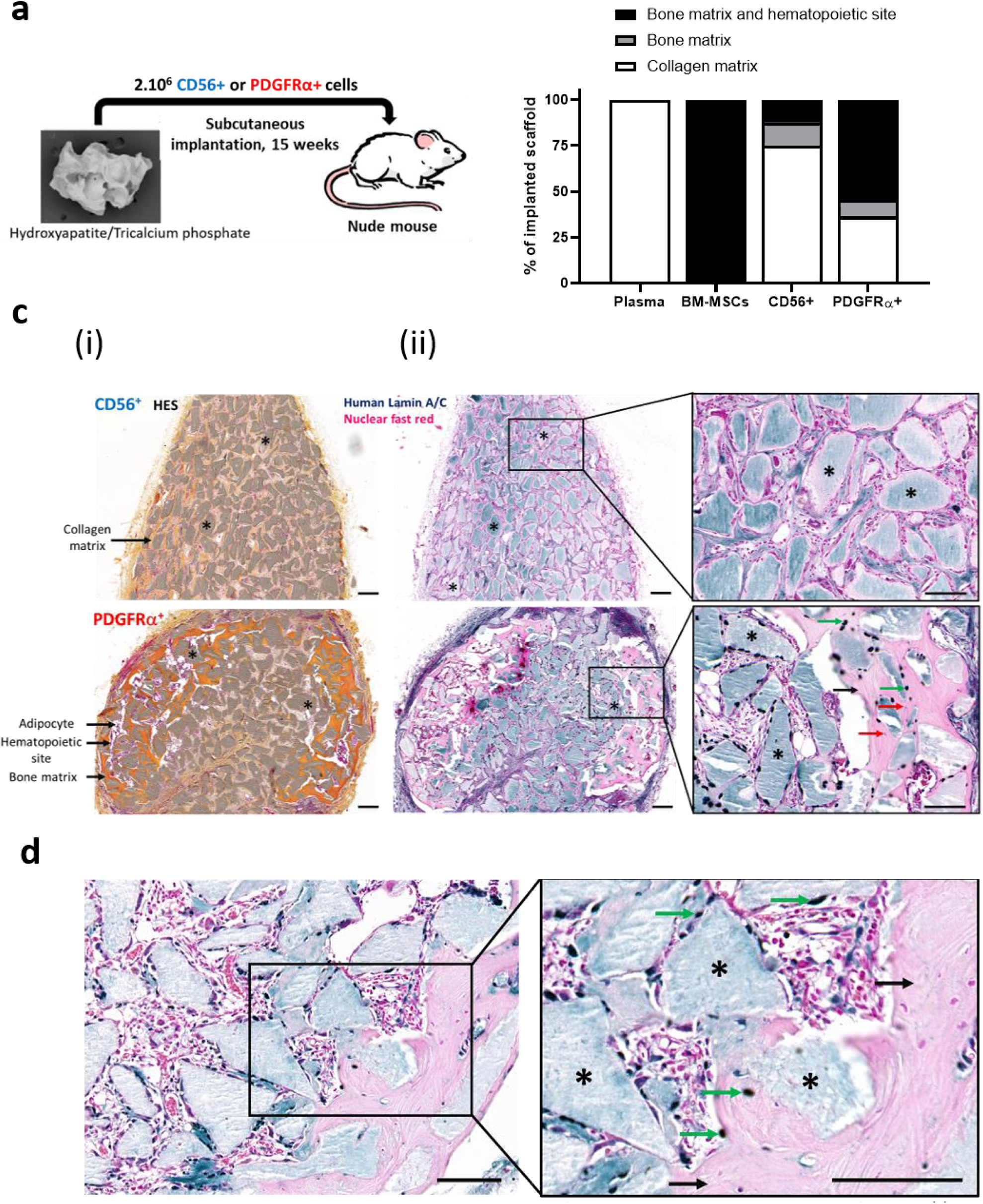
Human PDGFRα^+^ cells isolated from muscles surrounding NHO support *in vivo* bone formation. **a** Schematic representation of the *in vivo* osteogenic assay. Hydroxyapatite/calcium phosphate plasma scaffolds were seeded either with 2×10^6^ CD56^+^ cells, 2×10^6^ PDGFRα^+^ cells sorted from muscles surrounding NHO, 2×10^6^ human BM-MSCs or without cells (“plasma” negative control) and subcutaneously implanted in the back of nude mice for 15 weeks. **b** Percentage of scaffold containing either collagen matrix alone, bone matrix alone, or bone matrix associated with hematopoietic colonization. Human plasma (n=4 donors), BM-MSCs (n=6 donors), CD56^+^ cells (n=8; 4 donors, 2 implants/donor) and PDGFRα^+^ cells (n=11; 6 donors, 2 implants/donor for 5 donors and 1 implant for one donor). **c(i**)) Representative images of Hematoxylin-Eosin-Safran (HES) staining from CD56^+^ and PDGFRα^+^ cell seeded implant sections. Nuclei are stained in purple, cell cytoplasm in pink and collagen fibers in orange. * : hydroxyapatite. Magnification 10X; scale bar = 100µm. **c**(**ii**) Specific human Lamin A/C staining of a representative CD56^+^ and PDGFRα^+^ cell seeded implant sections. * : hydroxyapatite scaffold; black arrows: bone matrix; green arrow: human osteocytes; red arrows: mouse osteocytes. Magnification 10X and 20X; scale bar = 100µm. **d** Osterix/SP7 staining of a representative PDGFRα^+^ cell seeded implant section. * : hydroxyapatite scaffold; black arrows: bone matrix; green arrows: osterix^+^ osteoblasts. Magnification 20X and 40X; scale bar = 100µm.

## DISCUSSION

Although NHO develop mostly in peri-articular muscles in humans^1, 2^, their cellular origin remains unclear. Using *Pax7*^ZsG^ mice that specifically trace muscle SCs and their myogenic progenies, we clearly demonstrate that, NHO developing after SCI in injured muscles in mice are not derived from muscle SCs. We also reveal that NHO only develop in areas of the injured muscle where SCs fail to regenerate myofibers, with NHO only found in areas where ZsGreen-labeled SC-derived myocytes or myofibers are completely absent. Likewise, CD56^+^ muscle progenitors sorted from the muscles surrounding NHO from SCI patients, do not effectively generate ectopic bones when transplanted into permissive immunodeficient mice. Therefore, although NHO almost always develop in muscles, they are not the product of muscle SC trans-differentiation. In contrast, we find in *Prrx1*^ZsG^ mice focal accumulation and osteoblastic differentiation of *Prrx1*-expressing PDGFRα^+^ mesenchymal progenitor cells amongst areas of NHO development. This suggests that the SCI perturbs the normal or early proliferation followed by apoptosis of FAPs in the injured muscle with instead continued proliferation and absence of apoptosis of FAPs. High magnification of NHO sections in *Prrx1*^ZsG^ mice also reveal that osteocalcin^+^ osteoblasts and osteogenic progenitors contained in the developing NHO are labeled by ZsGreen, thus derived from *Prrx1*-expressing mesenchymal progenitor cells that share the same antigenic profile as muscle FAPs. Likewise, over half of PDGFRα^+^ mesenchymal progenitors sorted from the muscles surrounding NHO from SCI patients, effectively form ectopic bones when transplanted into permissive immunodeficient mice. One out 8 preparations of CD56^+^ muscle progenitor cells from these NHO biopsies could form ectopic bones with hematopoietic marrow when implanted in mice. Although it remains unclear whether this was caused by contaminating mesenchymal cells in the sorted population, our results suggest that in both mice and humans, NHO are derived from mesenchymal progenitor cells and not from myogenic progenitor cells such as SCs.

The *Pax7*Cre^ERT2^ model we employed has been shown by many groups to specifically target and trace SCs in muscles following injury^40, 50, 51^ which validates our use of *Pax7*^ZsG^ in our study. On the other hand, while there is no marker that is exclusively restricted to muscle FAPs, it is well known that *Prrx1*Cre (often called Prx1-Cre) targets and labels mesenchymal stromal cells in the developing limb buds as well their progenies in adults^41^. *Prrx1*Cre mice have been successfully used to target mesenchymal progenitor cells in adult bone marrow^52^, and mesenchymal progenitor cell-derived chondroprogenitors, osteoprogenitors, osteoprogenitors in bones, ligaments, tendons^53^ as well as adipocytes^54^, all of which are of mesenchymal origin^55^. Therefore, the mesenchymal specificity of the *Prrx1*Cre strain is well established. Herein, we show that in the skeletal muscle *Prrx1*Cre exclusively labels mesenchymal cells with the typical FAP phenotype (expressing CD34, Sca-1 and PDGFRα antigens at the cell surface, but not ITGA7 which is specific of SCs in the muscle^25^) both before or after muscle injury. Furthermore, these cells are reticulated and scattered along myofibers and do not form myofibers in the regenerating muscle, consistent with the known function of FAPs in coordinating muscle repair rather than forming new myocytes^25–29^. Therefore, although *Prrx1*Cre targets many other mesenchymal cells in various tissues of the body, it exclusively labels FAPs in the non-injured skeletal muscle. From these considerations, we conclude that our lineage-tracing models establish that the cells-of-origin which form NHO following a SCI are mesenchymal progenitor cells, not muscle satellite cells.

Other studies using models of FOP^32, 34^, calcific tendinitis and HO subsequent to tenectomy^44^ suggest that circulating mesenchymal progenitors contribute to HO formation. Therefore, we explored whether *Prrx1*-expressing cells were circulating following SCI or muscle injury using *Prrx1*^ZsG^ mice. As parabiosis experiments are banned in Australia for ethical reasons, we instead tracked these cells by flow cytometry of the peripheral blood. No cells expressing high levels of ZsGreen or with a FAP phenotype were detected in the blood of *Prrx1*^ZsG^ mice at multiple timepoints following SCI and muscle injury whereas these ZsGreen^high^ cells were very abundant in both naïve and injured muscles. Therefore, considering the high abundance of *Prrx1*-expressing FAPs in the muscle itself and their extreme rarity in the blood, it is very unlikely that NHO following SCI are derived from circulating mesenchymal cells. Although our SCI-induced NHO model does not involve BMP signaling and endochondral ossification, our result is consistent with a report showing that in model of HO induced by implantation of BMP-2 Matrigel, HO are not derived from circulating mesenchymal progenitors but from local FAPs^35^.

As lineage tracing experiments are not possible in humans, we attempted to validate our results from mice with NHO biopsies resected from SCI and traumatic brain injured patients. We isolated progenitor cell populations from skeletal muscle residues surrounding these resected NHO. Because of the small amount of muscle tissue on the surgical residues, adherent cells were amplified before and after sorting, possibly resulting in slight phenotypic changes. We sorted CD56^+^ cells known as myogenic progenitor cells^49^. Although they expressed CD90, CD105 and CD73 in culture, they were negative for PDGFRα and expressed key myogenic regulatory transcription factors *MY5* and *MYOD1*. Conversely, PDGFRα^+^ cells had the classical mesenchymal cell phenotype, were CD56^−^ and did not express myogenic regulatory factors *MY5* or *MYOD1*. Therefore, human muscle CD56^+^ and PDGFRα^+^ populations correspond to the *Pax7*- and *Prrx1*-expressing SCs and FAPs in mice. In our *in vitro* osteogenic assay, PDGFRα^+^ cells always had a higher osteogenic potential than CD56^+^ muscle progenitors, especially in the presence of pro-inflammatory cytokine OSM. PDGFRα^+^ cells were also more efficient in developing ectopic bones with human-derived osteocytes and osteoblasts and hematopoietic marrow when subcutaneously implanted in a conductive biomaterial in immunodeficient mice. Our findings are consistent with a previous report that showed that although both populations displayed osteogenic potential in vitro, PDGFRα^+^ cells had a far superior ability to form ectopic bones when transplanted on a supportive scaffold into immunodeficient mice as previously reported^56^. Taken together, these results strongly support the mesenchymal and non-myogenic origin of NHO in SCI patients.

Together, our lineage-tracing experiments suggest that NHO development subsequent to SCI is a pathology of the injured muscle, where muscle regeneration is deregulated with failure of muscle SCs to regenerate myocytes. We find that after a SCI, fewer FAPs undergo apoptosis at day 3 following muscle injury while FAPs continue to proliferate and incorporate BrdU even between days 12-14 post injury. This suggest that subsequent to a SCI, extensive and uncontrolled FAP survival, proliferation, differentiation results in the formation of extensive fibrotic areas in which osteogenic differentiation leads to heterotopic bone formation rather than muscle repair. This suggests that a severe CNS trauma such as SCI can reprogram FAPs in injured muscle whereby the apoptotic process that takes place in proliferative FAPs to reduce their numbers back to baseline levels after muscle injury^26, 27, 29^, is impaired in the context of a SCI. As a consequence, proliferating FAPs fail to apoptose and continue to accumulate in the injured muscle, which becomes fibrotic, followed by osteogenic differentiation of these FAPs into osteoblast-like cells leading to development of heterotopic bone tissue within injured muscles. From this perspective, we identified that SCI causes the upregulation of PDGFRα selectively at the surface of muscle FAPs independently of the muscle injury itself, with no such effect on SCs. This is consistent with the observation that sciatic nerve transection also upregulates PDGFRα expression in the denervated hindlimb muscle^28^ resulting in FAP accumulation and muscle fibrosis^28, 57, 58^. While heterotopic ossifications were not reported in these CDTX-injured denervated hindlimbs^57, 58^, we have shown that in the the presence of a SCI, denervation further increases NHO volumes^59^. Therefore our data suggest that the SCI reprograms FAPs in the skeletal muscles resulting in increased PDGFRα expression, reduced apoptosis and persistant proliferation in the injured muscles. As PDGF and its receptors are prime survival and proliferative signals in mesenchymal cells, this may contribute to NHO development. This will need to be further demonstrated in future work and it will be of interest to identify which factors (neural or systemic) cause PDGFRα up-regulation on FAPs in response to SCI.

In conclusion, our lineage tracing experiments in mice and xenotransplantation of human muscle cells associated with NHO reveal that NHO is a pathology of the injured muscle in which the SCI reprograms FAPs in the damaged muscle with fibrotic hyperproliferation and osteogenic differentiation instead of apoptosis. Importantly there is no trans-differentiation of muscle satellite cells. These our findings clarify the question of the cells-of-origin of NHO demonstrating for the first time that severe CNS traumas can reprogram FAPs in skeletal muscles. This new knowledge may provide new therapeutic opportunities to reduce NHO development in victims of severe CNS traumas.

## MATERIALS AND METHODS

### Animal ethics and sources

All experimental procedures in mice were approved by the Health Sciences Animal Ethics Committee of The University of Queensland and followed the Australian Code of Practice for the Care and Use of Animals for Scientific Purposes. In vivo osteogenic assays in nude mice were approved by the French Institutional Animal Care and Use Committee CAPSUD/N°26 under the ethics approval #9516/2017040715214163. C57BL/6 mice were obtained from Animal Resource Centre (Perth, Australia).

### Lineage tracing animal models

B6.Cg-*Pax7^tm1(Cre/ERT2)Gaka^*/J, B6.Cg-Tg(*Prrx1-cre*)1Cjt/J and B6.Cg-*Gt(ROSA)26Sor^tm6(CAG-ZsGreen1)Hze^*/J were purchased from the Jackson laboratory. B6.Cg-*Gt(ROSA)26Sor^tm6(CAG-ZsGreen1)Hze^*/J reporter strain (or ROSA26-LoxP-STOP-loxP-ZsGreen) has a CAG promoter-driven floxed STOP codon cassette ZsGreen cassette knocked into the *ROSA26* gene trap locus and therefore ZsGreen is only expressed once recombined by Cre recombinase. To specifically trace SCs, we selected B6.Cg-Pax7^tm1(Cre/ERT2)Gaka^/J (in brief *Pax7*-CreERT2) mice in which Cre recombinase is fused with a mutant estrogen receptor (CreERT2) fusion protein sequence that makes it tamoxifen-inducible and knocked-in together with an intra-ribosomal entry site downstream of the stop codon of the *Pax7* gene which is specifically expressed in SCs^40^. *Pax7*^ZsG^ mice were generated by crossing Pax7-CreERT2 to ROSA26-LoxP-STOP-loxP-ZsGreen mice (Fig. 1a). Primers and PCR conditions for genotyping are detailed in Supplementary table 2. CreERT2 nuclear translocation and subsequent ZsGreen expression by excision of the floxed STOP cassette in *Pax7*^ZsG^ mice was induced by daily oral gavage with tamoxifen 125 mg/kg for 4 days followed by a 2-week rest prior to surgery or CDTX intramuscular injection. To trace FAPs, we selected B6.Cg-Tg(Prrx1-cre)1Cjt/J transgenic mice (in brief *Prrx1*-Cre), in which Cre recombinase expression is controlled by the *Prrx1* promoter/enhancer specifically expressed in mesenchymal stem and progenitor cells. Prrx1^ZsG^ mice were generated by crossing *Prrx1*-Cre males to *ROSA26*-LoxP-STOP-loxP-ZsGreen female mice. Genotype of *Prrx1*-ZsGreen mice were confirmed by PCR as recommended by Jackson Laboratory.

### SCI-induced NHO mouse model

To induce NHO formation, 5-8 week old female mice underwent spinal cord transaction surgery (SCI) at T11-13 or control sham surgery (surgical incisions were only created on skin and muscle without damaging spinal cord) followed by an intramuscular injection (i.m.) of 0.32mg/kg purified Naja pallida CDTX (Latoxan, France) or equal volume of sterile phosphate buffered saline (PBS)^20^. Mice were euthanized by CO^2^ asphyxiation at the indicated time points. Hamstring muscle samples were processed for flow cytometry or histological analysis stated below.

### Flow cytometry

Single cells from hamstring muscles were isolated using a skeletal muscle dissociation kit (Miltenyi Biotech) and GentleMACS Dissociator tissue homogenizer (Miltenyi Biotec, Macquarie Park, Australia) as previously described^20^. Blood samples were taken by terminal cardiac puncture 1,2,3 and 7 days post-surgery in heparinized tubes under anesthesia (2-3% isoflurane). Red blood cells were lysed in 150mM NH^4^Cl, 10mM NaHCO^3^, 1 mM EDTA, pH=7.4 buffer with mixing and remaining cells were washed again in PBS containing 10% new-born calf serum (NCS), 2mM EDTA prior to staining.

To characterize ZsGreen^+^ SC-derived cells and ZsGreen^+^ FAP-derived cells, single cell suspensions were stained with monoclonal antibodies for surface markers: biotinylated anti-lineage (CD3ε, CD11b, B220, Gr1, Ter119) and streptavidin-APCCy7 CD45-BV785, CD31-BV421, CD34-e660, anti-Sca1-PECy7, anti-PDGFRα-BB700, anti-ITGA7-PE and Fixable Viability Stain 700 (FVS700). For PDGFRα expression analysis in C57BL/6 mice, cells were stained with surface markers: anti-Ter119-FITC, CD45-BV785, CD31-BV421, CD34-e660, anti-Sca1-PECy7, anti-PDGFRα-PE and FVS700. For mesenchymal stem cell mobilization, cells were stained with anti-lineage (CD3, B220, Ter119)-PerCpCy5.5, CD45-BV785, CD11b-BV510, F4/80-PE, CD31-BV421, CD34-e660, Sca1-PECy7 (antibodies detailed in Supplementary table 3). Flow analysis was performed using CytoFlex flow cytometer equipped with 405 nm, 488 nm, 561 nm and 640 nm solid state lasers (Beckman Coulter). Data analysis was performed using FlowJo v10.6.1 software following compensation with single color controls.

### MicroCT analyses

NHO volumes were measured in vivo in the Inveon PET-CT multimodality system (Siemens Medical Solutions Inc.) while mice were anesthetized with 2% isoflurane oxygen mixture. The parameters used were as follow: 360° rotation, 180 projections, 80 kV voltage, 500 μA current, and effective pixel size 36μm. After 3D image reconstruction, NHO volume was quantified in the Inveon Research Workplace (Siemens Medical Solutions Inc.) as previously described^20, 43^.

### Histological analyses of mouse muscles

Dissected muscle was fixed in 4% paraformaldehyde for 24 hours followed by decalcification in 14% EDTA for 4 days. Solution was replaced with 10, 20 and 30% sucrose/PBS every 24 hours. Muscles were mounted in optimum cutting temperature compound (OCT) and frozen in isopentane on liquid nitrogen and stored at −80°C. Muscle samples were sectioned at 8µm using Leica Cryostat CM1950. Every 8^th^ section was stained with hematoxylin & eosin and examined for the presence of heterotopic ossification (HO). Slides consecutive to those with confirmed HO were subsequently subject to immunofluorescent staining. After antigen retrieval with mild Proteinase K digestion (50ug/ml Proteinase K (Ambion) for 2min), sections were incubated for 10 min in PBS-Triton 0.5% and subsequently blocked for 1hr with 10% FBS/10% NGS in 0.1% Tween20 Tris-HCl buffered saline (TBS-Tween 0.1%). Sections were incubated with primary antibody (Collagen Type I (US Biological C7510-13), osteocalcin (EnzoLife Sciences ALX-210-333) or rabbit IgG control (ThermoFisher 31235)) diluted to 1ug/ml in TBS-Tween 0.1% for 1 hr at room temperature. Sections were washed with TBS-Tween 0.1%, incubated with biotin-labelled goat-anti-rabbit IgG secondary antibody (Vector Labs) for 20min, washed and incubated with AlexaFluor647 labelled streptavidin (ThermoFisher) for 20min. After counterstaining with 6-diamidino-2-phenylindole (DAPI) at 0.5ug/ml in TBS-Tween 0.1%, sections were mounted with ProLong™ Gold Antifade Mountant mounting medium (ThermoFisher). Images of histological stains were captured at 20X magnification using an Olympus VS120 Slide scanner Microscope. Fluorescent images were captured on a Perkin Elmer Vectra III Spectral Scanner Microscope and spectrally unmixed using InForm software (Perkin Elmer) or an Olympus BX63 Upright Epifluorescence microscope. The number of osteocalcin^+^ NHO nodules were counted from both *Pax7*^ZsG^ (n=4) and *Prrx1*^ZsG^ (n=3) mice. Two sections for the same mice (>120 µm apart) were stained, quantified and recorded in relation to the presence or absence of ZsGreen^+^ cells.

### Inhibition of BMP-2 signaling by LDN-193189 *in vitro*

Bone marrow mesenchymal cells^21^ were seed in a 96 well plate (1.5×10^3^/well) and maintained in αMEM + 20% FCS + 1% PSG till reaching confluence. Medium was then replaced with osteogenic medium (αMEM + 20% FCS + 1% PSG containing β-glycerophosphate (10mM), phosphor-ascorbic Acid (200µM), CaCl^2^ (2mM) and dexamethasone (0.2µM)) and recombinant human BMP-2 (100ng/ml) with additional LDN-193189 at 10, 100, 1000nM or equivalent volume of dimethyl sulfoxide vehicle. Medium were refreshed every 3 days. Mineralization on day7 was quantified using Alizarin red staining^21^. In brief, cells were washed with PBS and fixed in 4% paraformaldehyde for 30 minutes. After rising off fixative, cells were air-dried overnight and stained with 1% Alizarin red S solution at RT on shaker for 5-10 minutes then washed with PBS and air-dried overnight. Staining was dissolved in 10% cetylpyridinium Chloride (CPC) containing sodium phosphate (10mM) at RT on shaker for 15 minutes and transferred to a new 96 well plate to measure observance at 562nm using Thermofisher Multiskan reader.

### Mouse muscle RNA extraction and qRT-PCR

Female C57BL/6 mice underwent SCI or Sham surgery followed by i.m. injections of CDTX or PBS. Four days after surgery, hamstring muscle samples were harvested and snap frozen in liquid nitrogen. Frozen muscle was homogenized in Trizol with IKA T10 basic homogenizer on ice for 30 seconds for 3 times with 10 second intervals followed by incubation at RT for 30 mins. Tissue debris was removed after centrifuging at 12,000xg and supernatant was transferred to fresh tubes and mixed with 1/5 volume of chloroform. After spinning down at 12,000xg, 4 C for 15 mins, aqueous phase was transferred to fresh tube and RNA was precipitation by isopropanol, washed in 75% ethanol, air-dried and dissolved in RNAse/DNase free water. RNA was quantified using nanodrop (ThermoScientific). cDNA was synthesized using SeniFast cDNA synthesis kit following manual instructed by company. The mRNA expression was analyzed using single-step reverse transcription quantitative real-time polymerase chain reaction (RT-qPCR) Taqman system. Reaction was prepared following instruction of TaqMan™ fast PCR Master Mix and *Bmp2, Bmp4, Bmp7* and *Rps20* assay (Supplementary table 4). Reactions were read using Applied Biosystems Viia7 Real-time PCR system. Ct values were normalized by the expression of house-keeping gene *Rps20* and presented as ratio to house-keeping gene.

### In vivo LDN-193189 treatments

C57BL/6 mice underwent SCI and CDTX intramuscular injection as indicated above on day 0. Mice were treated with vehicle (6% DMSO in PBS) or LDN-193189 (30mg/kg) via intraperitoneal (i.p.) injection twice daily from day 0-14 or day14-21 a dosage which is effective in reducing HO volumes in mouse model of FOP driven by *ACVR1*^Q207D^ missense mutation^45^. Animals were randomly allocated to vehicle or treatment groups after surgery (while animals were still under anesthesia) but prior to treatment commencements. Therefore, grouping was not affected by recovery of surgery.

### In vivo FAP apoptosis measurement

Mice underwent SCI or sham surgeries and CDTX injection and harvested 3 days post-surgery. Muscle cells were isolated as described above followed by surface marker staining: biotinylated anti-lineage (CD3ε, CD5, B220, CD11b, Gr1, Ter119), CD45-BV785, CD31-BV421, CD34-e660, Sca1-PECy7 and anti-ITGA7-PE antibodies followed by streptavidin (SAV)-BUV395. After surface maker staining, cells were washed in 1X PBS and then annexin binding buffer (10mM Hepes pH=7.4, 150mM NaCl, 5mM MgCl^2^, 5mM KCl, 1.8mM CaCl^2^). Next cells were stained with Annexin V-FITC (1/50 dilution) in annexin binding buffer at RT for 20 mins. After washing off excess Annexin V, cells were resuspended in annexin binding buffer and add 1/5 volume of 7AAD provided in the kit. Flow cytometry analysis was performed using BD LSR Fortessa X20 cytometer equipped with 355 nm, 405 nm, 488 nm, 561 nm and 640 nm solid state lasers.

### In vivo FAP proliferation measurement by BrdU incorporation

To measure in vivo FAP proliferation during NHO development, mice underwent SCI or sham surgeries and CDTX injection. Two days prior to tissue harvest at day 14 post-surgery, mice were given drinking water containing 0.5mg/ml BrdU (Sigma-Aldrich cat#B5002) together with twice daily i.p. injection of BrdU for the last 2 days before tissue sampling. One mouse from SCI+CDTX group and one mouse from Sham+CDTX group were not treated with BrdU and used as negative control for BrdU staining.

Muscle cells were isolated as described above followed by surface marker staining: anti-lineage (CD3ε, CD5, B220, CD11b, Gr1, Ter119)-Pacific blue, CD45-BV785, CD31-PE, CD34-e660, anti-Sca1-PECy7. Cells were then fixed and permeabilized using BrdU flow kit (BD Pharmingen, cat#559619) following manufacturer’s instruction. Fixed cells were then treated with DNase I (13 µg / 1-2 x 10^6^ cells) in DNAse reaction buffer (10mM Tris-HCl, 0.5mM CaCl^2^, 2.5mM MgCl^2^ in PBS, pH7.4) at 37°C for 1 hour. BrdU incorporation in DNA was labeled with anti-BrdU-FITC monoclonal antibody in buffer containing 10 µg / mL blocking mouse IgG at RT for 1 hour. Cells were then washed and analyzed by flow analysis was performed using BD LSR Fortessa X20 cytometer.

### Isolation of human BM-MSCs, PDGFRα^+^ and CD56^+^ cells

All NHO and muscle samples were obtained with the informed consent of the patients and the approval from the people protection committee (CPP approval n°09025) and from the National Commission for Informatics and Liberties (CNIL approval n° Eyo1066211J). NHO biopsies were taken from the hip of 12 patients: 7 patients with traumatic brain injury (age: 28, 35, 35, 48, 52, 63, 71), 3 patients with SCI (age 26, 33, 65) and 2 patients with stroke (age 47, 72).

Muscle surrounding NHOs were collected from NHO surgical waste following their excision from patients with brain injuries, spinal cord injuries or strokes. NHO resection surgeries were performed at Raymond Poincaré Hospital (Garches, France). Muscle fragments were minced using scalpel and small scissors, placed in a 50 ml Falcon tube and incubated in 1.5 mg/ml pronase (#10165921001 Sigma-Aldrich) in α-MEM, 45min in a 37°C water bath. After addition of α-MEM supplemented with 15% FCS and 1% antibiotics, the cell suspension was filtered through a 100 µm cell strainer followed by a 40 µm cell strainer (BD Falcon). Isolated muscle progenitor cells (MPCs) were maintained in α-MEM supplemented with 15% FCS, 1% antibiotics and 10 ng/ml basic fibroblast growth factor (bFGF) (R&D Systems). Human MPCs were trypsinized and incubated 30 min with biotinylated anti-human PDGFRα (R&D system) and CD56-PE (clone B159, BD Pharmigen) monoclonal antibodies in PBS 2% FCS, 2mM EDTA. Cells were washed and incubated 30 min with Streptavidin-APC/Cy7 and the viability dye 7-AAD (Sony). Cells were washed and filtered through a 30µm cell strainer (Sysmex) and sorted using a FACSAria III SORP sorter (BD Biosciences). PDGFRα^+^ and CD56^+^ cells were seeded at 3,000 per cm^2^ in α-MEM supplemented with 20% FCS, 1% antibiotics and 10 ng/ml bFGF (R&D Systems).

BM-MSCs were isolated by plastic adhesion and expanded up to 70-80% confluence in α-MEM supplemented with 10% FCS and 1% antibiotics. Cells were subsequently seeded at 4,000 per cm^2^.

### Flow cytometry phenotyping of human PDGFRα^+^ and CD56^+^ cells

PDGFRα^+^ and CD56^+^ cell suspensions (passages 4 and 5) were stained with monoclonal antibodies for surface markers CD31-PE, CD34-APC, CD45-APC, CD73-PE, CD90-APC, CD105-PE (Supplementary table 3). Flow analysis was performed using CytoFlex flow cytometer (Beckman Coulter). Data analysis was performed using FlowJo v10.6.1 software following compensation with single color controls.

### qRT-PCR of human PDGFRα^+^ and CD56^+^ cells

PDGFRα^+^ and CD56^+^ cells (passages 3 to 5) were lysed in Qiazol (Qiagen) followed by chloroform/isopropanol total RNA extraction. Reverse transcription was performed using Reverse Transcriptase Core Kit (Eurogentec). qRT-PCR were performed on a LightCycler 480 instrument (Roche) using QuantiTect SYBR Green RT-PCR Kit and QuantiTect Primers (Qiagen). Three housekeeping gene mRNAs (*GAPDH*, *PPIA*, *HPRT*) were selected based on their expression stability using geNorm analysis. *MYF5* and *MYOD1* quantification were performed as the geometric mean of the quantifications obtained with each reference RNA.

### In vitro osteogenic differentiation and quantification of mineralization of human cells

PDGFRα^+^ and CD56^+^ cells were seeded in 24-well plates at 3,000 per cm in α-MEM supplemented with 10% FCS and 1% antibiotics. Once cells adhered to the wells, medium was replaced by α-MEM supplemented with 10% FCS, 1% antibiotics and 0.052 µg/ml dexamethasone, 12.8 µg/ml ascorbic acid, and 2.15 mg/ml ß-glycerophosphate (Sigma-Aldrich) to induce osteogenic differentiation. Human recombinant OSM was added at 100 ng/ml (Miltenyi Biotec). Cells were cultured for 14 days at 37°C in 5% CO^2^ atmosphere, and medium was changed twice a week. Quantification of mineralization was performed using Alizarin Red S staining. Cells were washed with PBS, fixed in 70% ethanol, quickly washed with distilled water and incubated 5 min in 20g/L Alizarin Red S, pH=4.2 (Sigma-Aldrich). Cells were washed with distilled water and dried. Alizarin Red S dye was extracted with 0.5N hydrochloric acid and 5% SDS and quantified by spectrophotometry at 405 nm.

### In vivo osteogenic assays with human cells

Subconfluent human BM-MSCs, PDGFRα^+^ and CD56^+^ cells (2-3 amplification passages) were collected for subcutaneous implantation into the flanks of 10 weeks old male nude mice (Janvier Labs). Implants were prepared by mixing 2.10^6^ cells to sterile 80–200 µm particles (60% hydroxyapatite/40% ß-tricalcium phosphate) (Graftys), 100 µl of human plasma in a 1ml syringe as described^60^. Control “plasma implants” did not contain cells. Ten microliters of 2% CaCl^2^ in water was added to induce coagulation. Fifteen weeks after implantation, animals were euthanized, scaffolds were collected and fixed in 4% paraformaldehyde overnight at 4°C. The following day, scaffolds were washed with PBS and decalcified in 20% EDTA pH=8 solution for 48 hr. Implants were washed with PBS and dehydrated with graded series of ethanol solutions prior to paraffin embedding (Thermo Scientific).

### Histology and human lamin A/C immunohistochemistry

Paraffin sections (4μm) were dried, deparaffinized, and stained with Hematoxylin Eosin Safran (HES) (Dako). For immunohistochemical staining, 4µm paraffin sections were incubated with antigen retrieval citrate-based solution pH=9 (Vector Laboratories) at 95°C for 15 min. Sections were incubated in Bloxall blocking solution (Vector Laboratories) for 10 min to inactivate endogenous peroxidase. FC receptor blocking reagent (Innovex biosciences) was then added for 45 min. All sections were incubated in PBS-FCS 10% for 30 min and anti-lamin A/C antibody (dilution 1/200, clone EPR4100, Abcam) diluted in PBS-FCS 5%-BSA 1.5% was then added for overnight incubation at 4°C. Sections were washed with PBS-Triton 0.1% and incubated in ImPRESS REAGENT anti-rabbit IgG (Vector Laboratories) for 30 min. Then, sections were washed with PBS-Triton 0.1% and incubated in HistoGreen substrate solution (Linaris) for 1 min. Counter coloration was performed using with Fast Nuclear Red (Vector Laboratories). All sections were analyzed using a Pannoramic Midi II slide scanner and Case Viewer software (3D HISTECH Ltd.).

### Statistics

Statistic differences were calculated using Mann-Whitney test or one-way ANOVA with Tukey’s multiple comparison test with Prism v7 or 8 software (GraphPad software, La Jolla, CA).51. Data visualisation in Fig. 4 was also performed using Rstudio v1.1456, tidyverse and ggplot2 packages.

## ACKNOWLEDGEMENTS

This work was partly supported by Project Grant 1101620 and Ideas Grant 1181053 from the National Health and Medical Research Council of Australia (NHMRC) and by award W81XWH-15-1-0606 from the Congressionally Approved Spinal Cord Injury Research Program of the US Department of Defense and by funds from the Mater Foundation. JPL is supported by Research Fellowship 1136130 from the NHMRC. This work was also partly funded by project grant 1101620 from the French Government Defense Procurement and Technology Agency (DGA – Direction Générale de l’Armement). The Translational Research Institute is partly funded by the Federal Government of Australia.

The authors greatly acknowledge Sabrina Soaves, Bastien Rival and Muriel Nivet for technical assistance. The authors would like to thank the technical assistance of “Cochin HistIM Facility” (Paris, France). They also highly acknowledge Pr Philippe Denormandie and Dr Laure Gatin, (Service de Chirurgie Orthopédique, Hôpital Raymond Poincaré, Garches, France) and Guillaume Genêt for providing and coordinating access to NHO and muscle surgical residues. The authors also acknowledge the scientific and technical assistance at the Translational Research Institute: The University of Queensland biological resources TRI facility, histology facility, flow cytometry facility, microscopy facility and the preclinical imaging facility, which is supported by Therapeutic Innovation Australia (TIA). TIA is supported by the Australian Government through the National Collaborative Research Infrastructure Strategy (NCRIS) program.

## AUTHOR CONTRIBUTIONS

JPL, SB and MCLBK conceived, designed and supervised the study. HWT, DG, SM, KA, FT, AA, WF, JG, MEG, DC, BN, MS and BJ conducted experiments, analyzed data and interpreted results. AP, MS, FG contributed to scientific discussion, interpretation and experimental design. HWT, DG, SM, KA and WF prepared figures for publication. HWT, DG, MCLBK, SB and JPL prepared the manuscript.

---

**Table S1.**
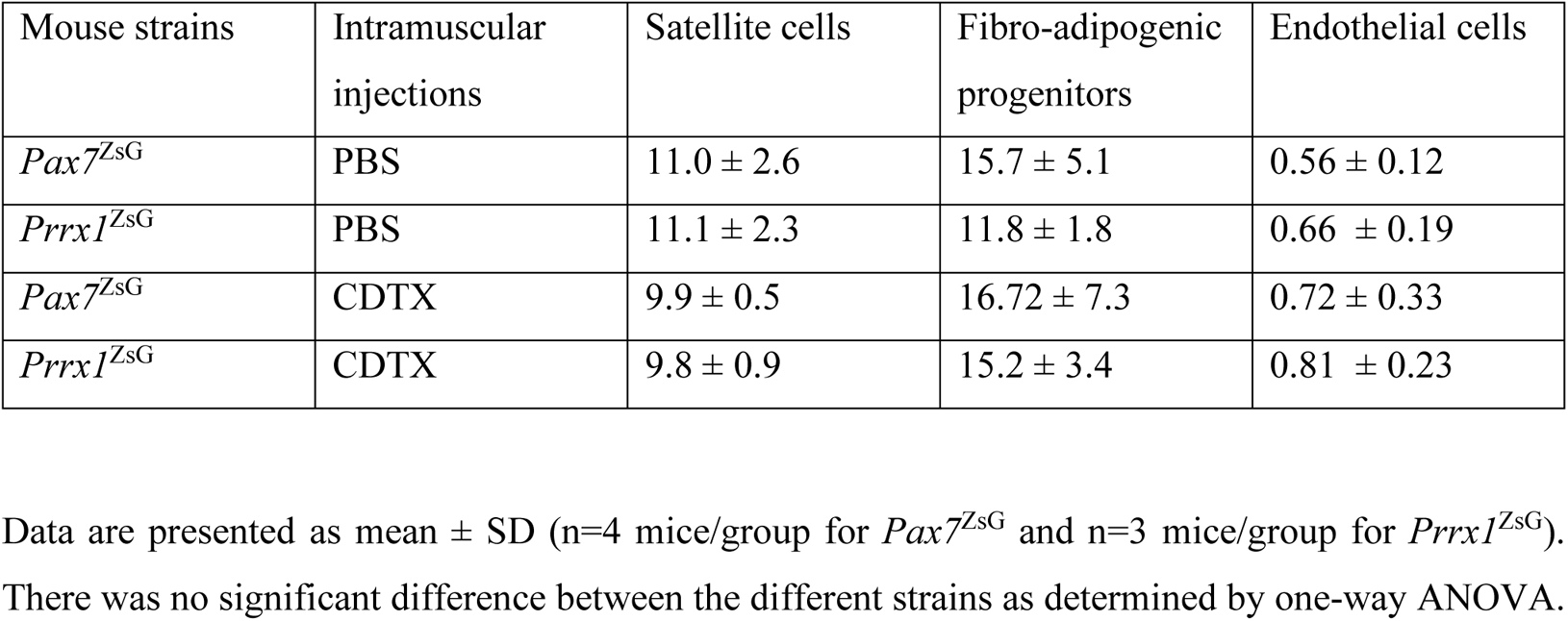
Percentage of phenotypic SCs, FAPs and endothelial cells in non-hematopoietic cells from the hamstring muscles of *Pax7*^ZsG^ and *Prrx1*^ZsG^ mice at 14 days post PBS or CDTX injection measured by flow cytometry.

**Table S2.**
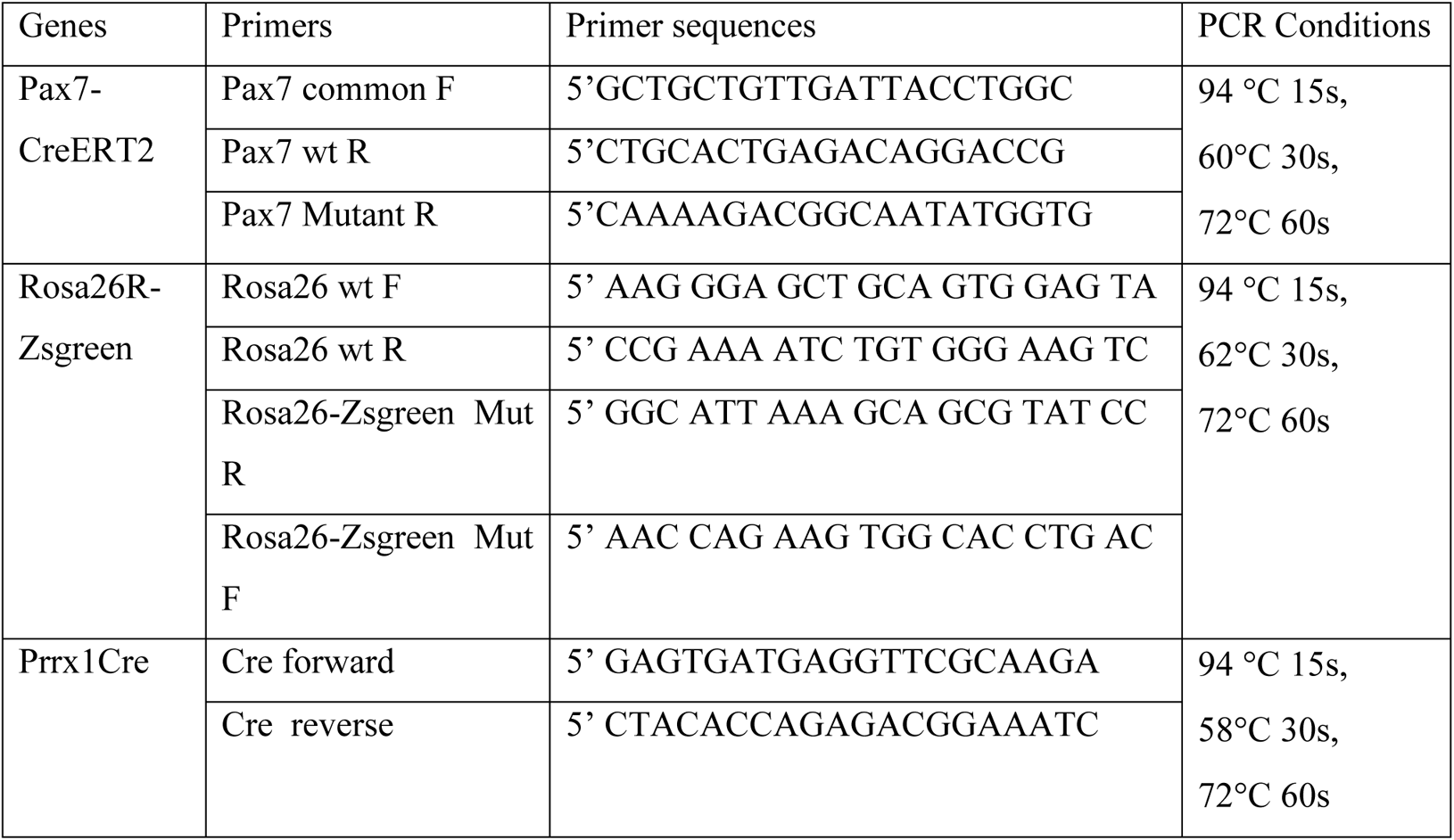
Genotyping PCR reagents and conditions

**Table S3.**
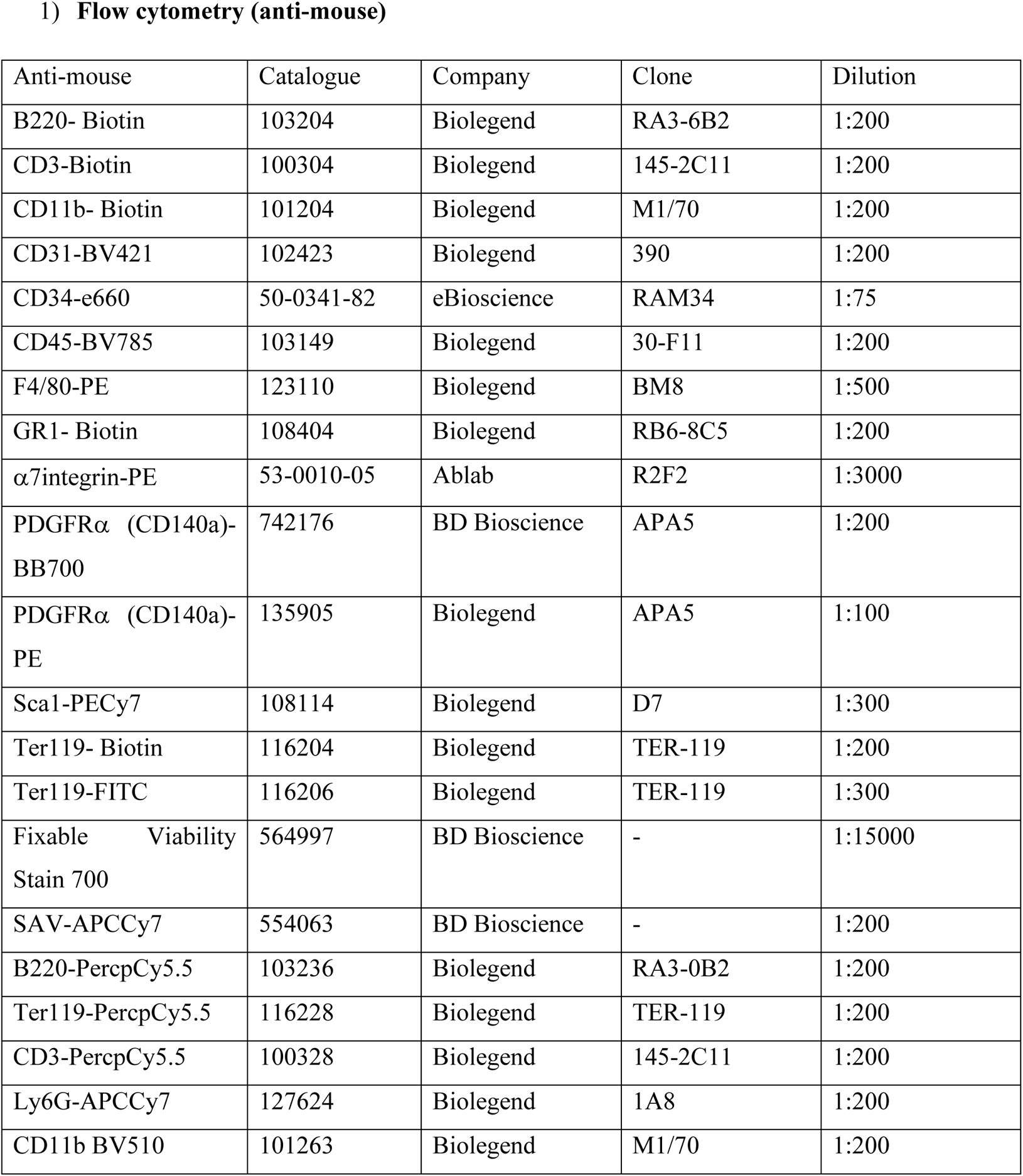
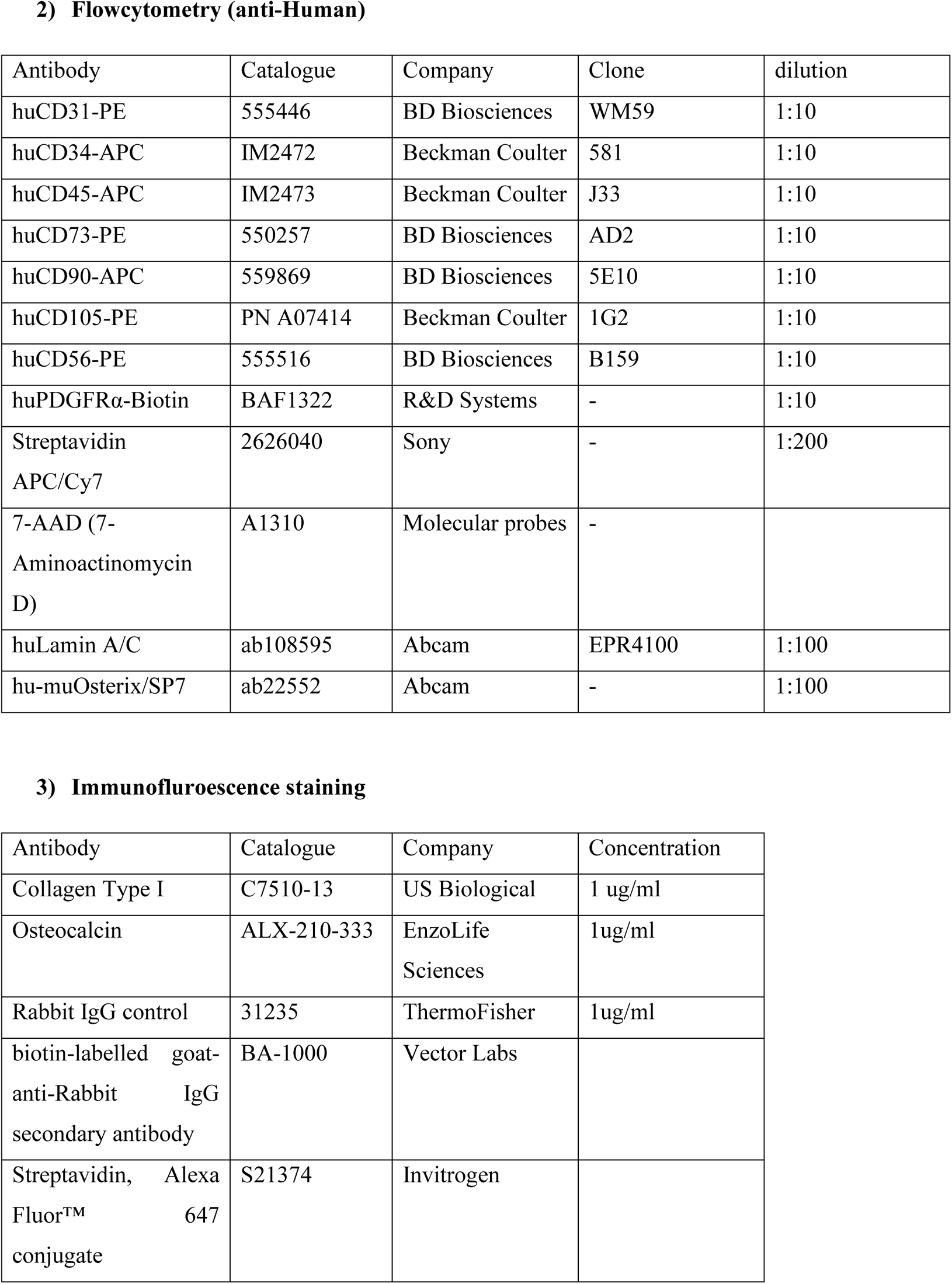
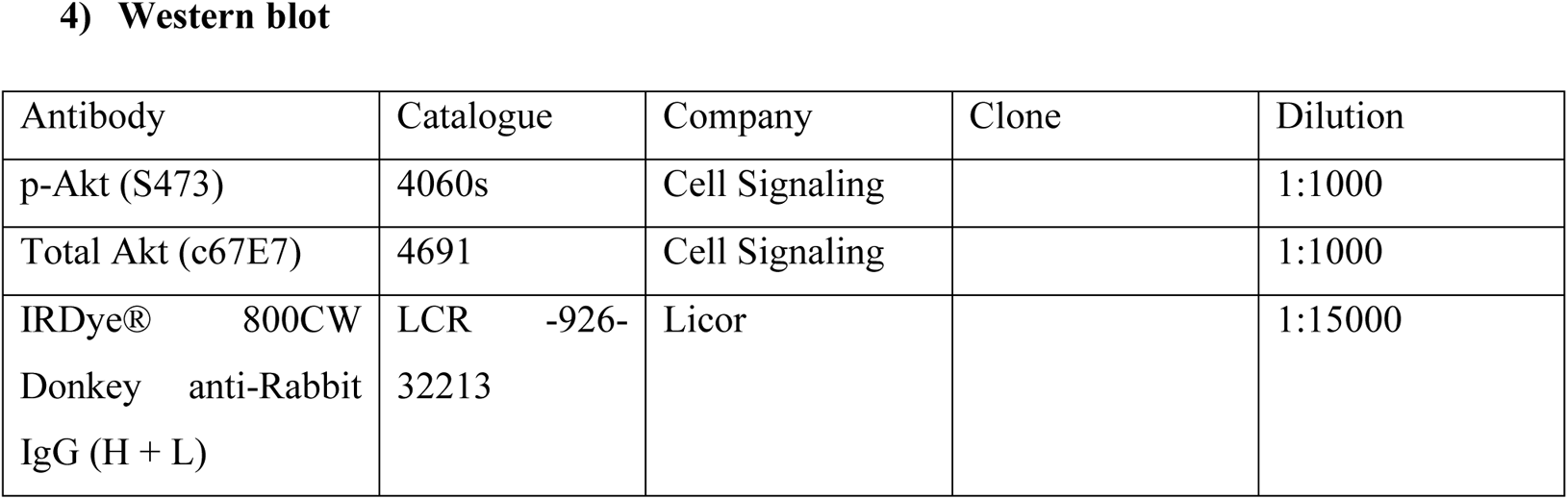
Antibodies used in the study.

**Supplemental Table 4:**
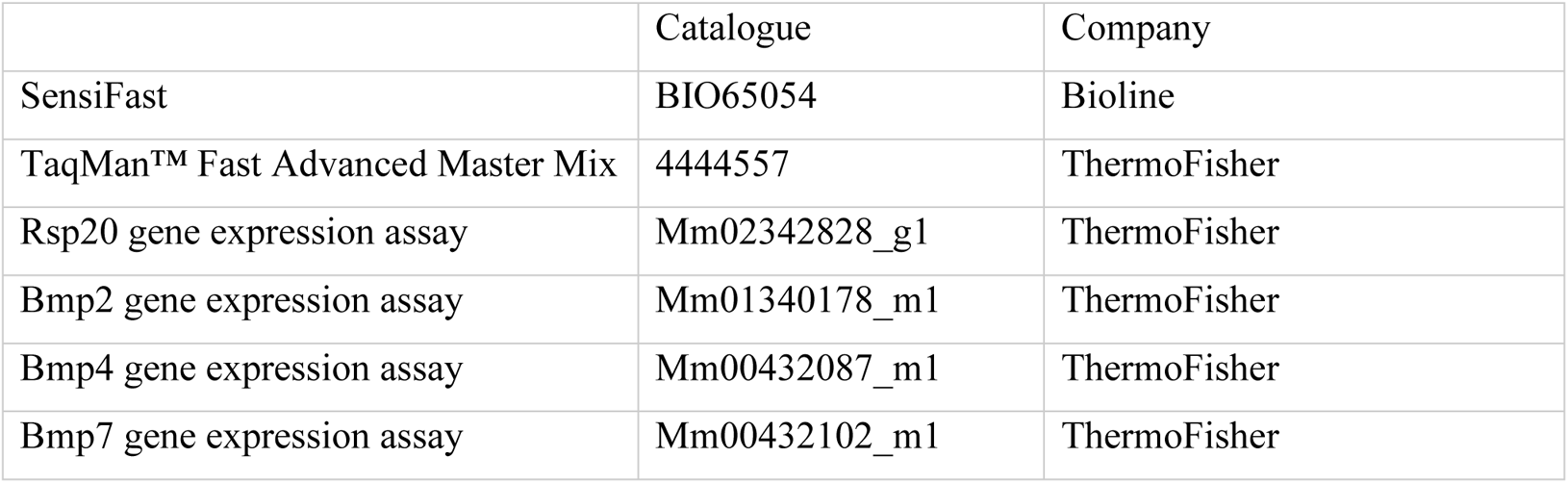
**qRT-PCR primer probe sets**

**Supplemental Table 5.**
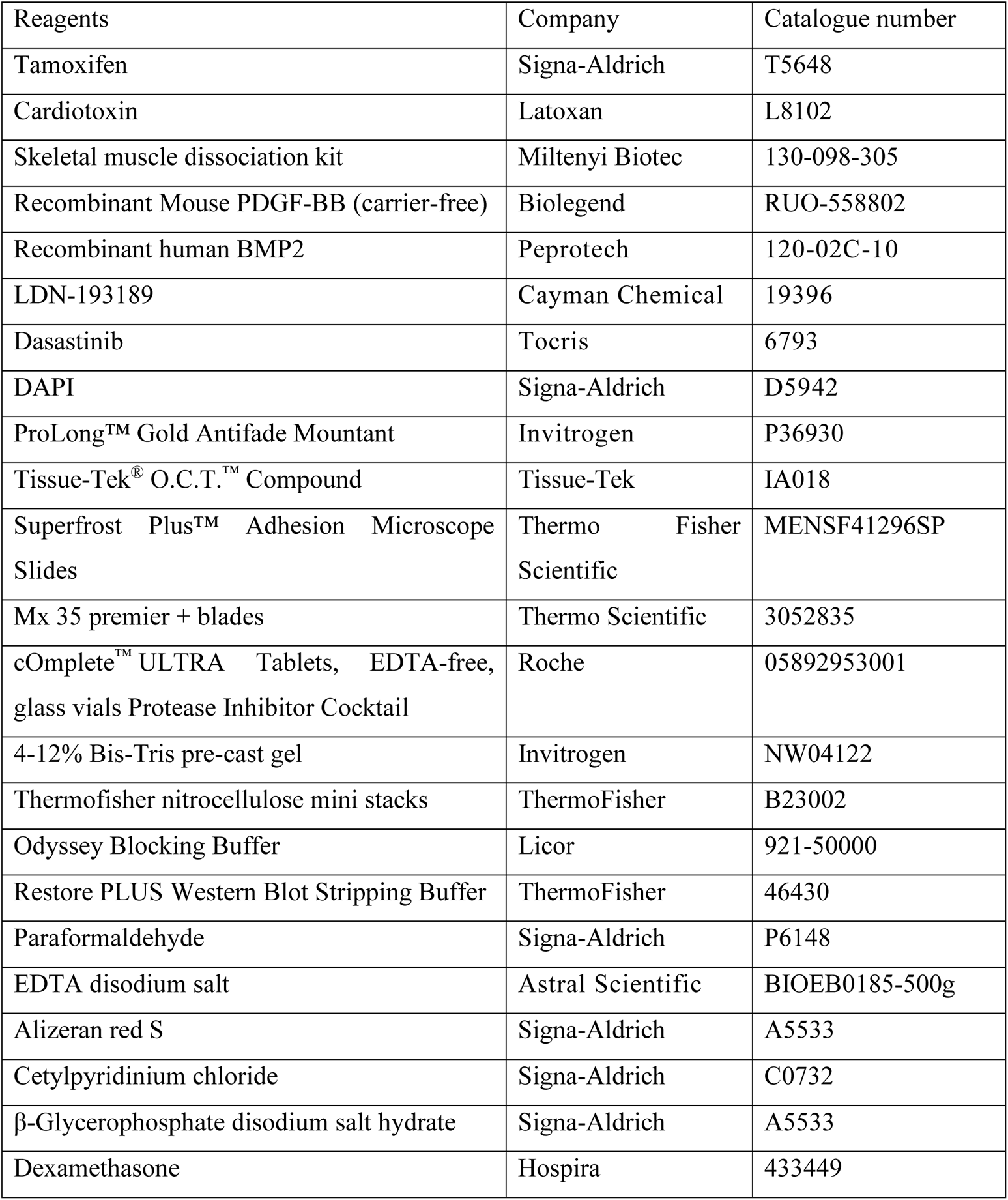
**Other reagents.**

**Fig. S1.**
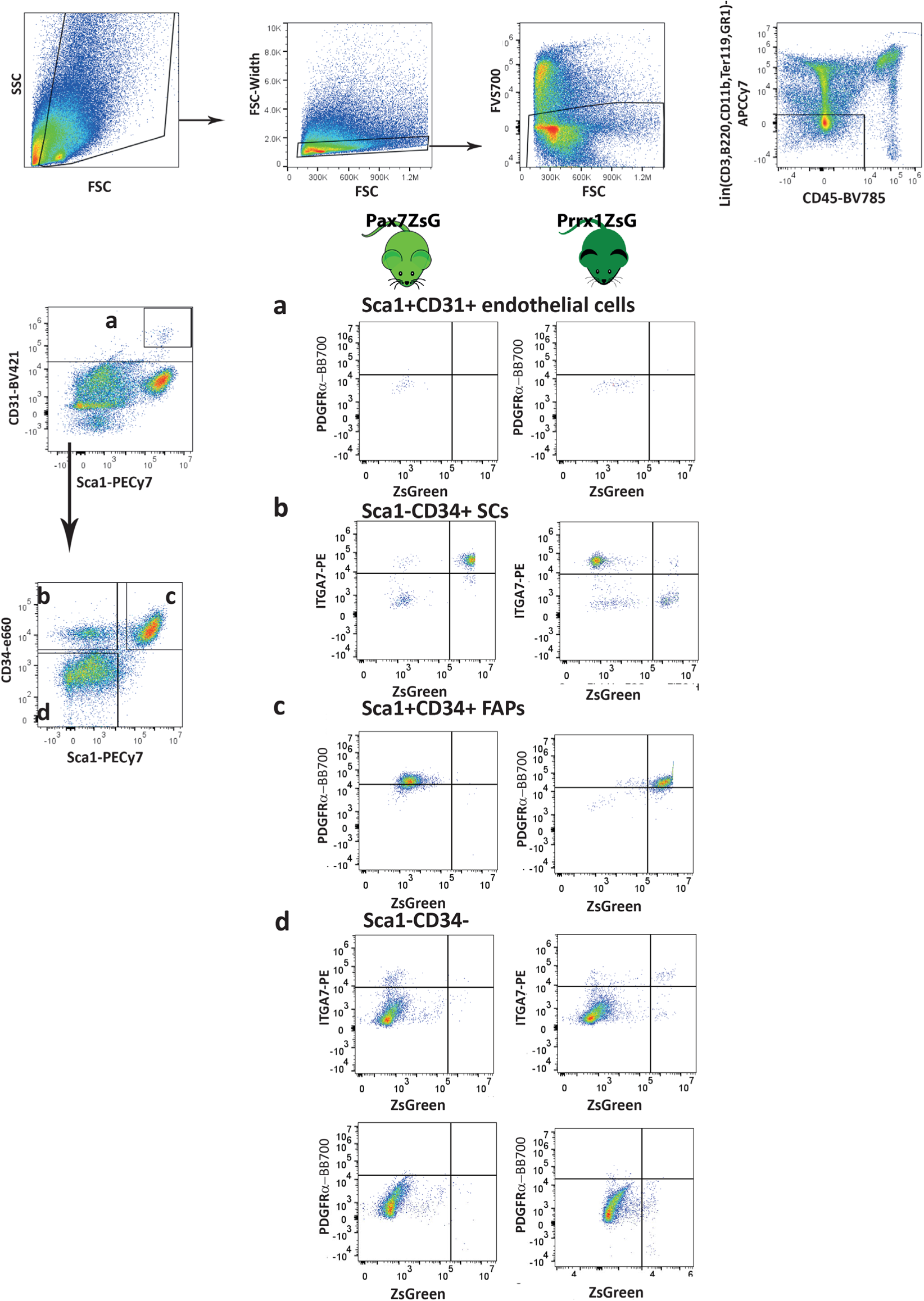
Flow cytometry gating strategy to identify SCs and FAPs. Representative dot-plots from PBS-injected hamstring muscles from *Pax7*^ZsG^ (n=4 /group) and *Prrx1*^ZsG^ (n=3 /group) mice from the same data sets as shown in Fig. 1. After gating on forward/side scatters and FVS700 dead cell exclusion, Lin^−^ CD45^−^ non-haematopoietic cells was gated for **a** Sca1^+^CD31^+^ endothelial cells, **b** CD31^+^ Sca1^−^ CD34^+^ SCs, **c** CD31^−^ Sca1^+^ CD34^+^ FAPs and **d** CD31^−^ Sca1^−^ CD34^−^ population. Expression of ZsGreen versus integrin α7 (ITGA7) and PDGFRα were further analyzed in *Pax7*^ZsG^ (left panels) and *Prrx1*^ZsG^ (right panels). **(A)** Sca1^+^ CD31^+^ endothelial cells were ZsGreen^−^ PDGFRα^−^ in both strains. **b** CD31^−^ Sca1^−^ CD34^+^ ITGA7^+^ SC are ZsGreen^+^ in *Pax7*^ZsG^ mice but ZsGreen^−^ in *Prrx1*^ZsG^ mice. **c** CD31^−^ Sca1^+^ CD34^+^ PDGFRα^+^ FAPs are ZsGreen^−^ in *Pax7*^ZsG^ mice but ZsGreen^+^ in *Prrx1*^ZsG^ mice. **d** CD31^−^ Sca1^−^ CD34^−^ population in Pax7^ZsG^ are ZsGreen^−^ while there are small populations of cells expressing ITGA7 but no expression of PDGFRα. On the other hand, in *Prrx1*^ZsG^ mice, a small population express ZsGreen and ITGA7 but not PDGFRα.

**Fig. S2.**
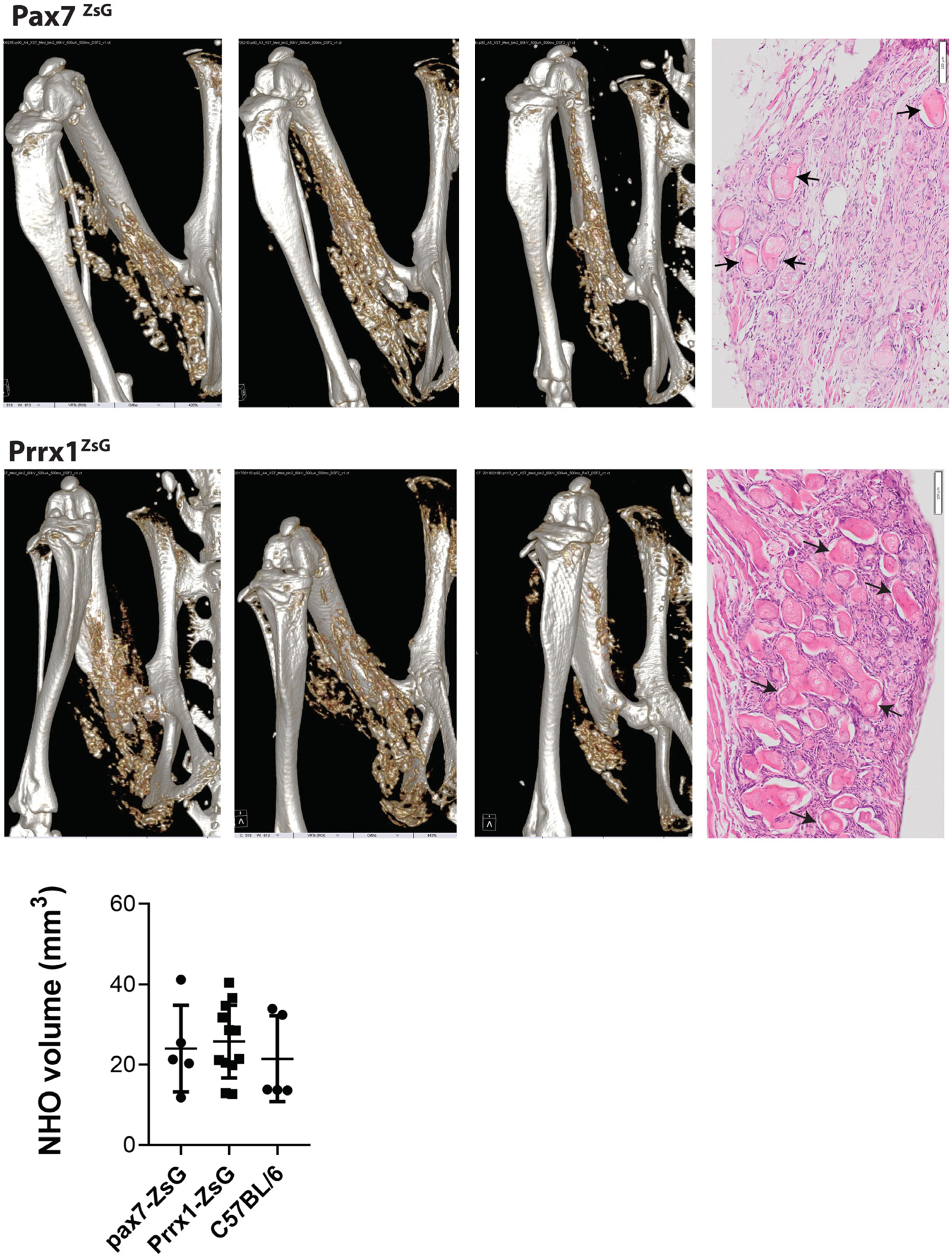
*Pax7*^ZsG^ and *Prrx1*^ZsG^ mice underwent SCI surgery and i.m. injection of CDTX. NHO development was quantified by CT on day7 as representative images. Each point indicates one mouse and data was presented as mean ±S D. Statistic difference was calculated by Kruskal-Wallis test. Muscle were harvested on day28 and stained with Hematoxylin and eosin. Arrows indicates small bone nodules within necrotic muscle tissues.

**Fig. S3.**
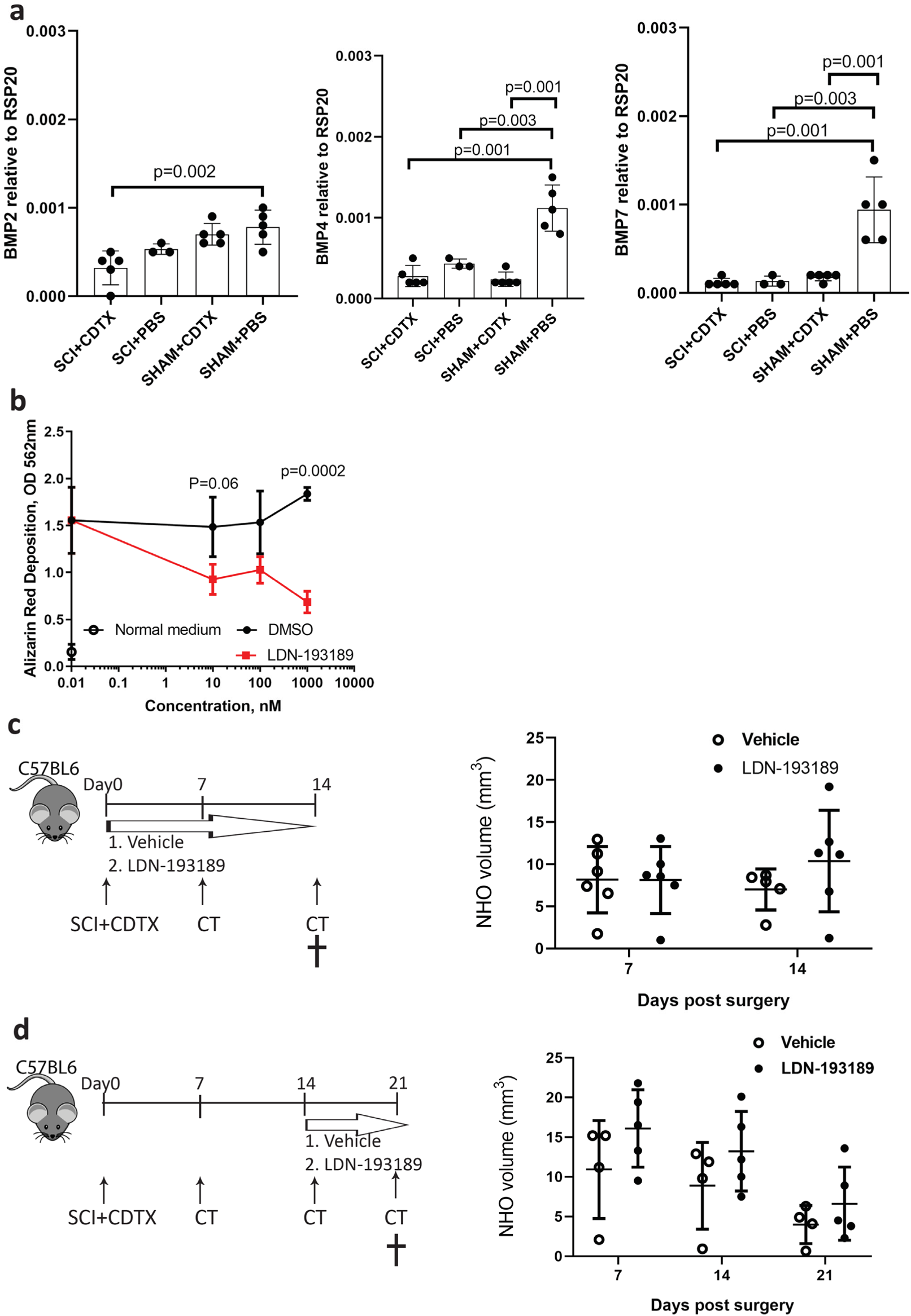
BMP signalling is not required in the initiation and development of NHO. **a** *Bmp2*, *Bmp4* and *Bmp7* mRNA expression in muscles taken from SCI+CDTX, SCI+PBS, Sham+CDTX and Sham+PBS 4 days post-surgery were analysed by qRT-PCR. Expression was normalised by housekeeping gene *Rsp20*. Each point indicates one mouse and data was presented as mean±SD. Statistic difference was calculated by one-way ANOVA with Sidak multiple comparison analysis. **b** Mouse bone marrow mesenchymal cells were cultured with normal medium (open circles), osteogenic medium with recombinant BMP-2 (100ng/ml), DMSO vehicle (black full circles) or LDN-193189 (red squares) at indicated concentration. Mineralisation was quantified on day7 using Alizarin red and absorbance was read at 562nm. Data was presented as mean ± SD (triplicate/treatment) and statistic difference was analysed by two-way ANOVA and Bonferroni’s multiple comparisons test. **c** Mice were treated with vehicle or LDN-193189 at 3 mg/kg twice daily by i.p. injection from day 0-14 and NHO were quantified by μCT on day7 and 14 (n=6 mice/group). **d** Mice were treated with vehicle or LDN-193189 (3 mg/kg twice daily) by i.p. injection from day 0-14 and NHO were quantified by μCT on day7, 14 and 21 (n=5 mice/group). Each point indicates one mouse and data was presented as mean ±S D. Statistic difference was calculated by Mann-Whitney test for comparison at each time point.

**Fig. S4.**
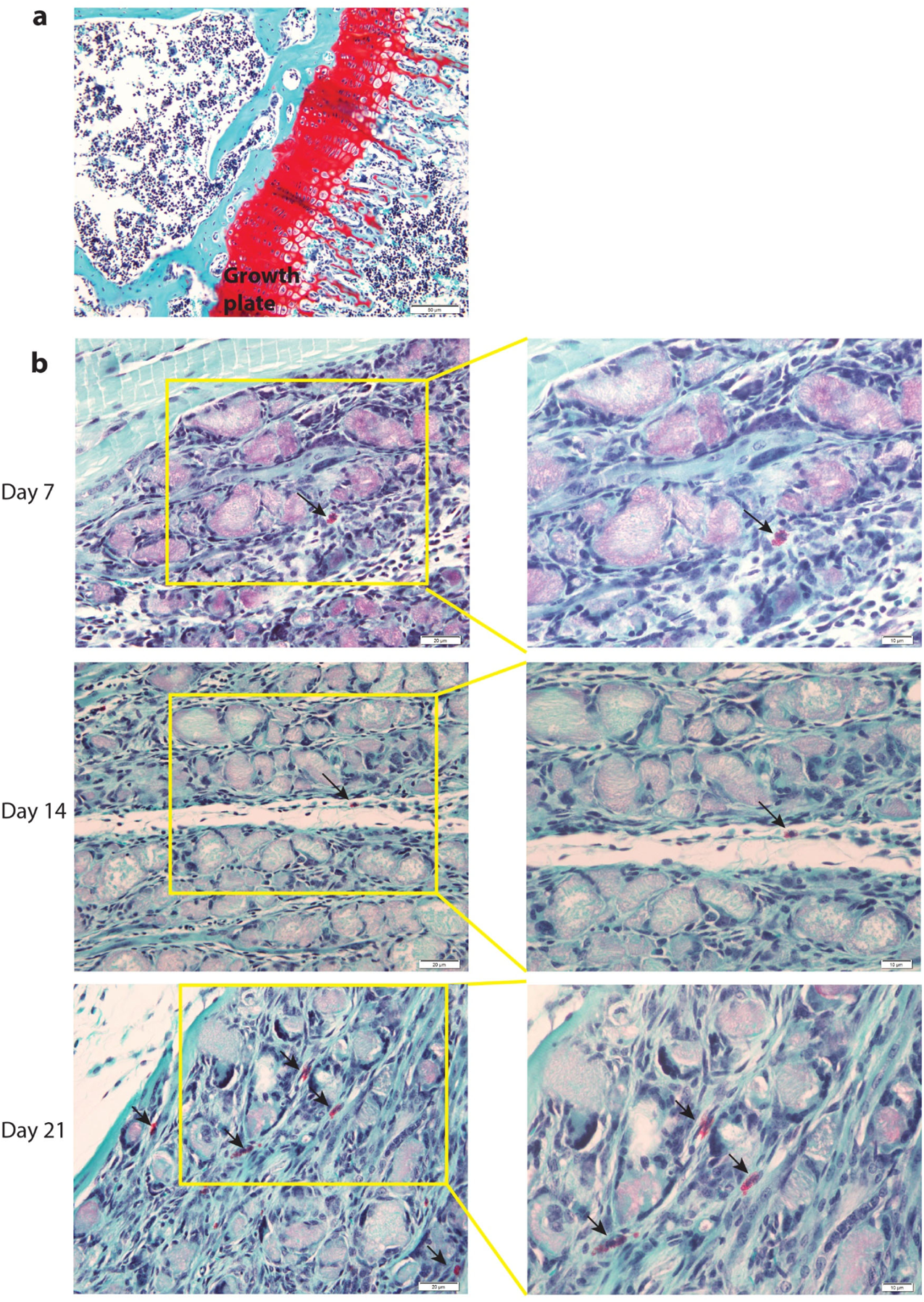
NHO do not develop via endochondral ossification following SCI in mice. Safranin O/fast green staining was performed on injured muscle samples harvested 7, 14 and 21 days post SCI+CDTX surgery to examine whether NHO formation involves endochondral ossification. **a** Proteoglycan in growth plate stained bright red as positive control. **b** Occasional mast cells (arrows) with characteristic basophilic granules scattering in the injured muscle samples across day 7, 14 and 21 stained with Safranin O. However, bright red chondrocytes or cartilage matrix could not be detected in NHO at any time-point (n=3-6 mice per time point). Figures in the right column are higher magnification of yellow boxes in the left column. Scale bar: a 50 µm, b left =20µm, right =10µm.

**Fig. S5.**
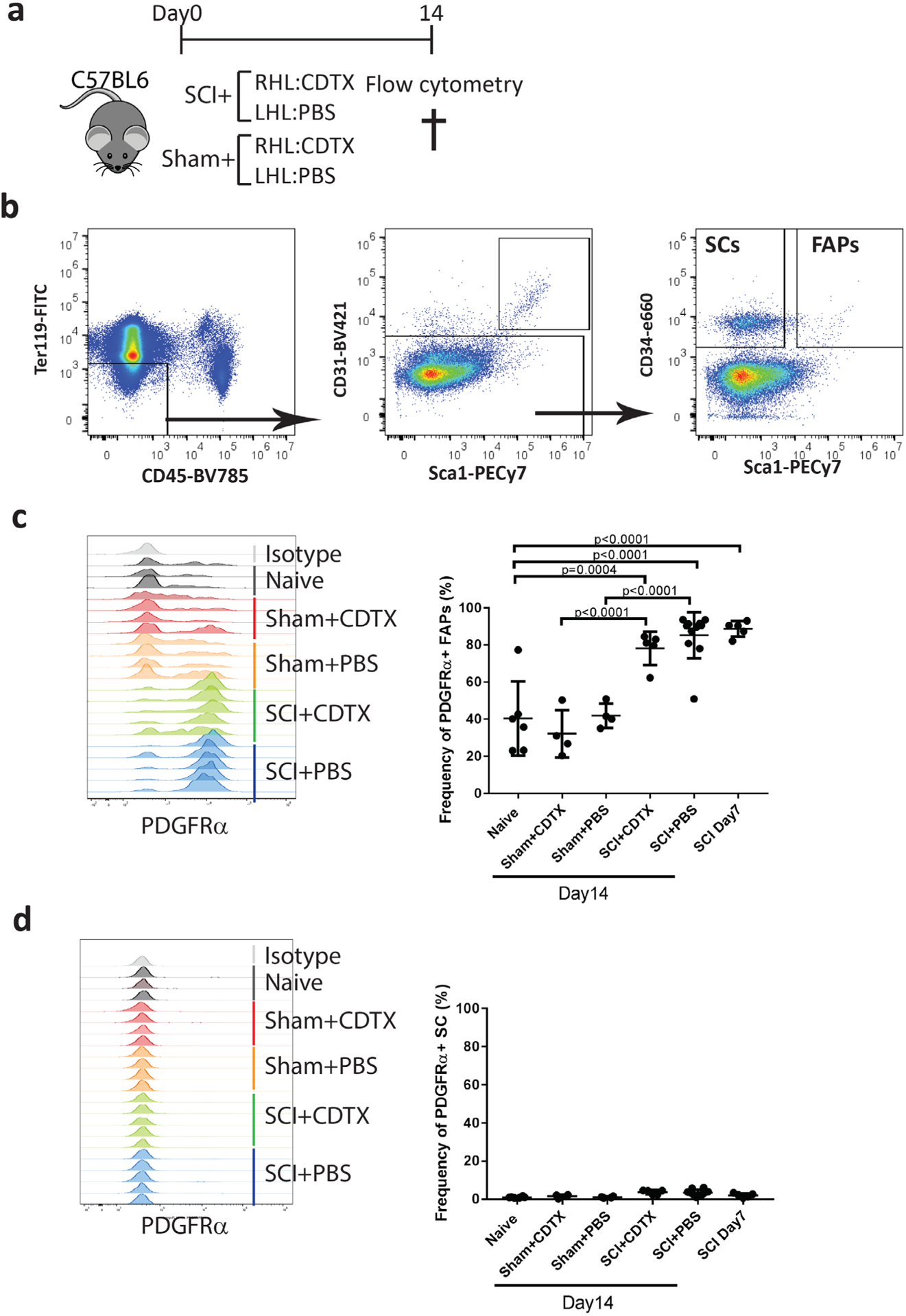
SCI upregulates PDGFRα expression on FAPs but not SCs. **a** C57BL/6 mice received SCI or Sham surgery plus intramuscular injection of CDTX in RHL and PBS in LHL. Hamstring muscle cells were subsequently isolated 7 or 14 days post-surgery or from naïve mice. **b** FAPs and SCs were gated from live CD45-Ter119-CD31-cells and separated according to their expression of Sca1 and CD34. Overlayed histograms of PDGFRα expression on **c** Sca1^+^ CD34^+^ FAPs and **d** Sca1^−^ CD34^+^ SCs isolated from naïve (grey), sham + CDTX (red), sham + PBS (orange), SCI + CDTX (green), and SCI + PBS (blue) groups (n=3-5mice/group). Frequency of PDGFRα expression in FAPs and SCs of is shown as mean ± SD. Each dot represents a separate mouse. Significance was calculated by one-way ANOVA with Tukey’s multiple comparison test.

**Fig. S6.**
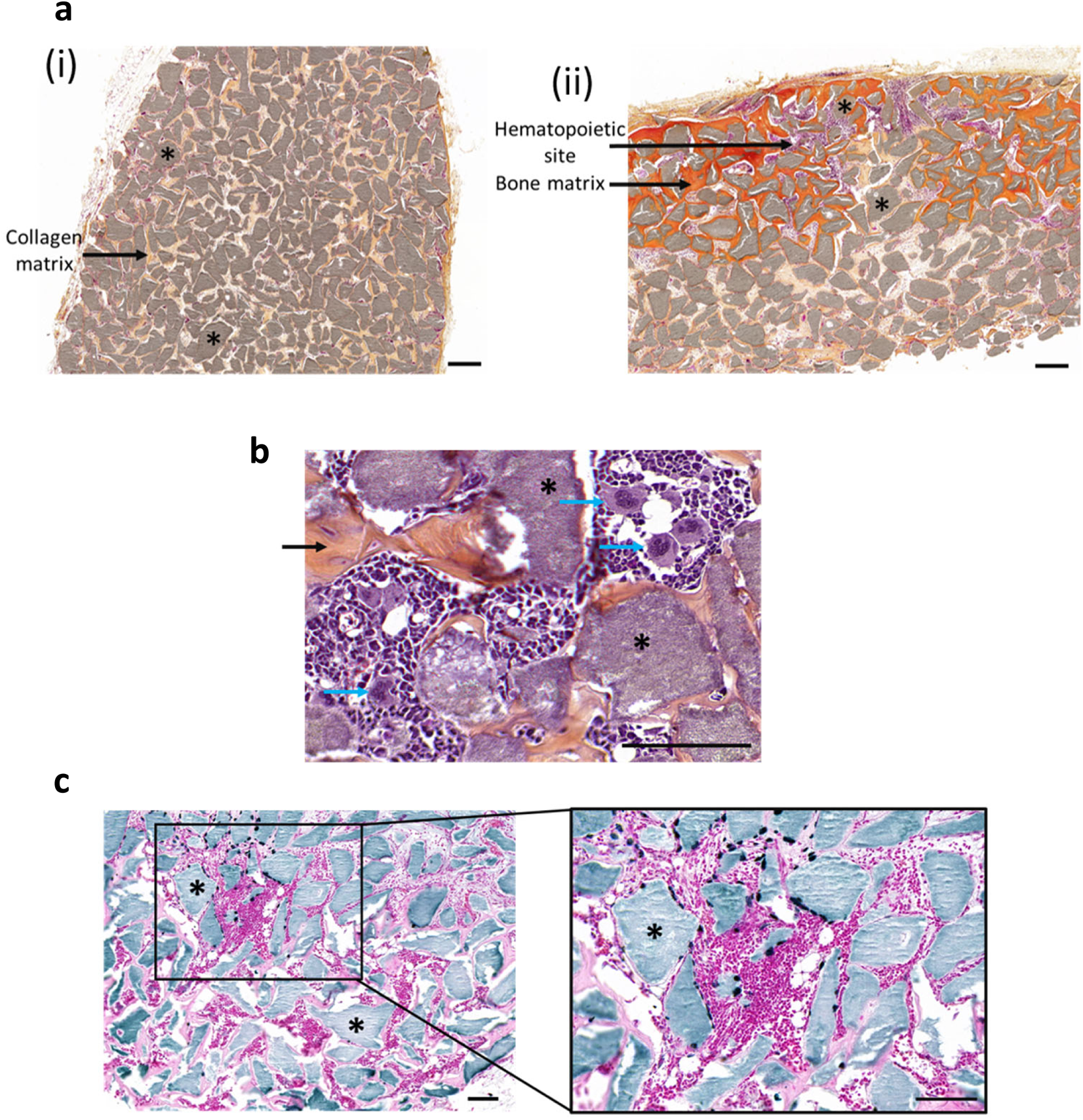
*In vivo* osteogenic assay: Histological and IHC characterization of implanted scaffolds. **a** Representative images of Hematoxylin-Eosin-Safran (HES) staining from (**i**) plasma implant section and (**ii**) BM-MSCs cells seeded implant sections. * : hydroxyapatite. Magnification 10X; scale bar = 100µm. **b** Representative images of Hematoxylin-Eosin-Safran (HES) staining from PDGFRα^+^ cell seeded implant section showing large mature megakaryocytes (blue arrows). * : hydroxyapatite scaffold; black arrow: bone matrix. Magnification 40X; scale bar = 100µm. **c** Specific human Lamin A/C staining of a representative PDGFRα^+^ cell seeded implant section. * : hydroxyapatite scaffold. Magnification 10X and 20X; scale bar = 100µm.

