## Supplementary tables for "Spinal cord injury reprograms muscle fibro-adipogenic progenitors to form heterotopic bones within muscles"

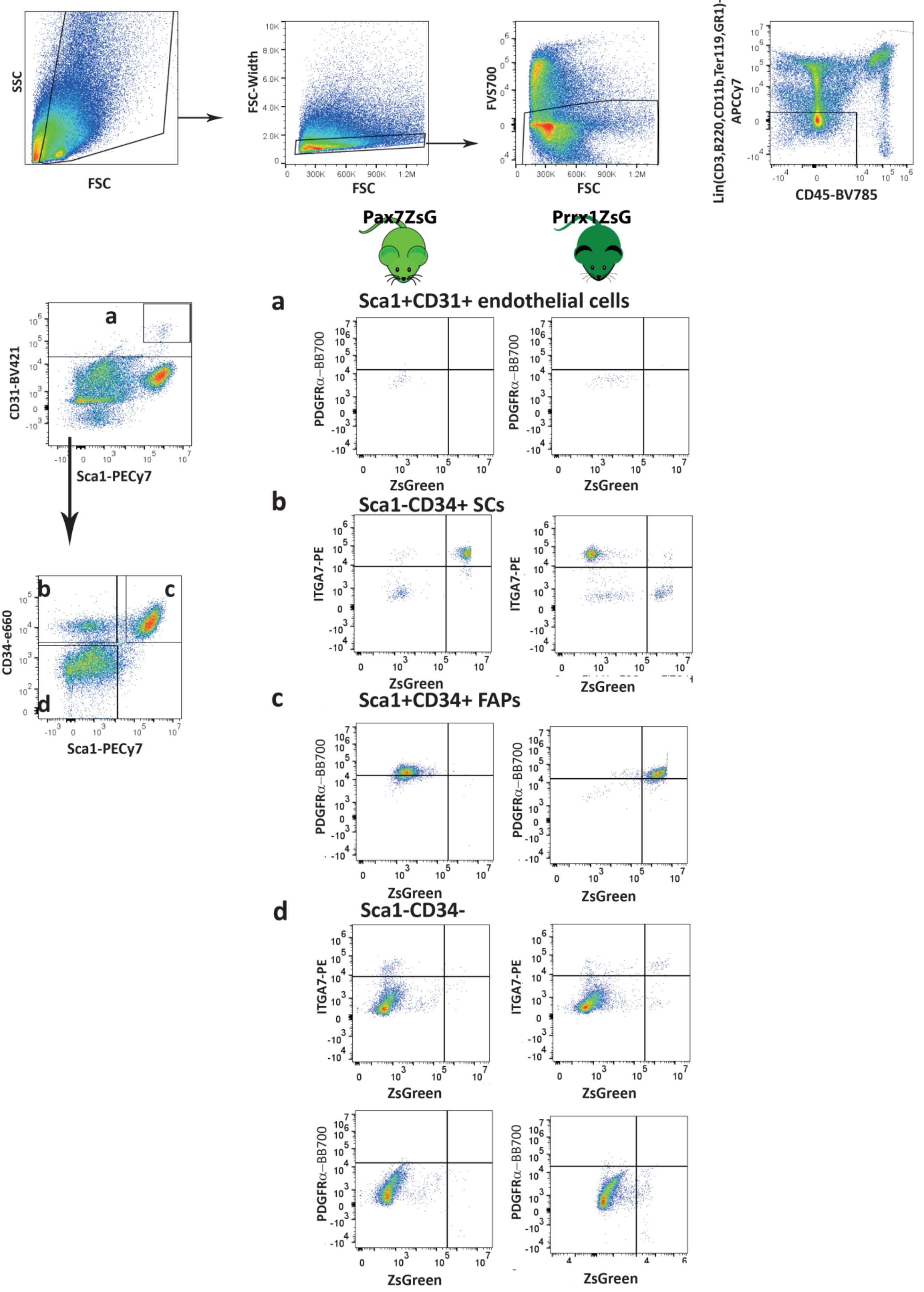

SSC
FSC
FSC-Width
FSC
FVS700
FSC
Lin(CD3,B220,CD11b,Ter119,GR1)-APCCy7
CD45-BV785
Pax7ZsG
Prrx1ZsG
CD31-BV421
Sca1-PECy7
a
CD34-e660
Sca1-PECy7
b
c
d
a
Sca1+CD31+ endothelial cells
PDGFRα–BB700
ZsGreen
b
Sca1-CD34+ SCs
ITGA7-PE
ZsGreen
c
Sca1+CD34+ FAPs
PDGFRα–BB700
ZsGreen
d
Sca1-CD34-
ITGA7-PE
ZsGreen
PDGFRα–BB700
ZsGreen


Figure S1

**Fig. S1**. Flow cytometry gating strategy to identify SCs and FAPs. Representative dot-plots from PBS-injected hamstring muscles from *Pax7*$^{ZsG}$ (n=4 /group) and *Prrx1*$^{ZsG}$ (n=3 /group) mice from the same data sets as shown in Fig. 1. After gating on forward/side scatters and FVS700 dead cell exclusion, Lin$^{-}$ CD45$^{-}$ non-haematopoietic cells was gated for **a** Sca1$^{+}$CD31$^{+}$ endothelial cells, **b** CD31$^{+}$ Sca1$^{-}$ CD34$^{+}$ SCs, **c** CD31$^{-}$ Sca1$^{+}$ CD34$^{+}$ FAPs and **d** CD31$^{-}$ Sca1$^{-}$ CD34$^{-}$ population. Expression of ZsGreen versus integrin α7 (ITGA7) and PDGFRα were further analyzed in *Pax7*$^{ZsG}$ (left panels) and *Prrx1*$^{ZsG}$ (right panels). **(A)** Sca1$^{+}$ CD31$^{+}$ endothelial cells were ZsGreen$^{-}$ PDGFRα$^{-}$ in both strains. **b** CD31$^{-}$ Sca1$^{-}$ CD34$^{+}$ ITGA7$^{+}$ SC are ZsGreen$^{+}$ in *Pax7*$^{ZsG}$ mice but ZsGreen$^{-}$ in *Prrx1*$^{ZsG}$ mice. **c** CD31$^{-}$ Sca1$^{+}$ CD34$^{+}$ PDGFRα$^{+}$ FAPs are ZsGreen$^{-}$ in *Pax7*$^{ZsG}$ mice but ZsGreen$^{+}$ in *Prrx1*$^{ZsG}$ mice. **d** CD31$^{-}$ Sca1$^{-}$ CD34$^{-}$ population in Pax7$^{ZsG}$ are ZsGreen$^{-}$ while there are small populations of cells expressing ITGA7 but no expression of PDGFRα. On the other hand, in *Prrx1*$^{ZsG}$ mice, a small population express ZsGreen and ITGA7 but not PDGFRα.

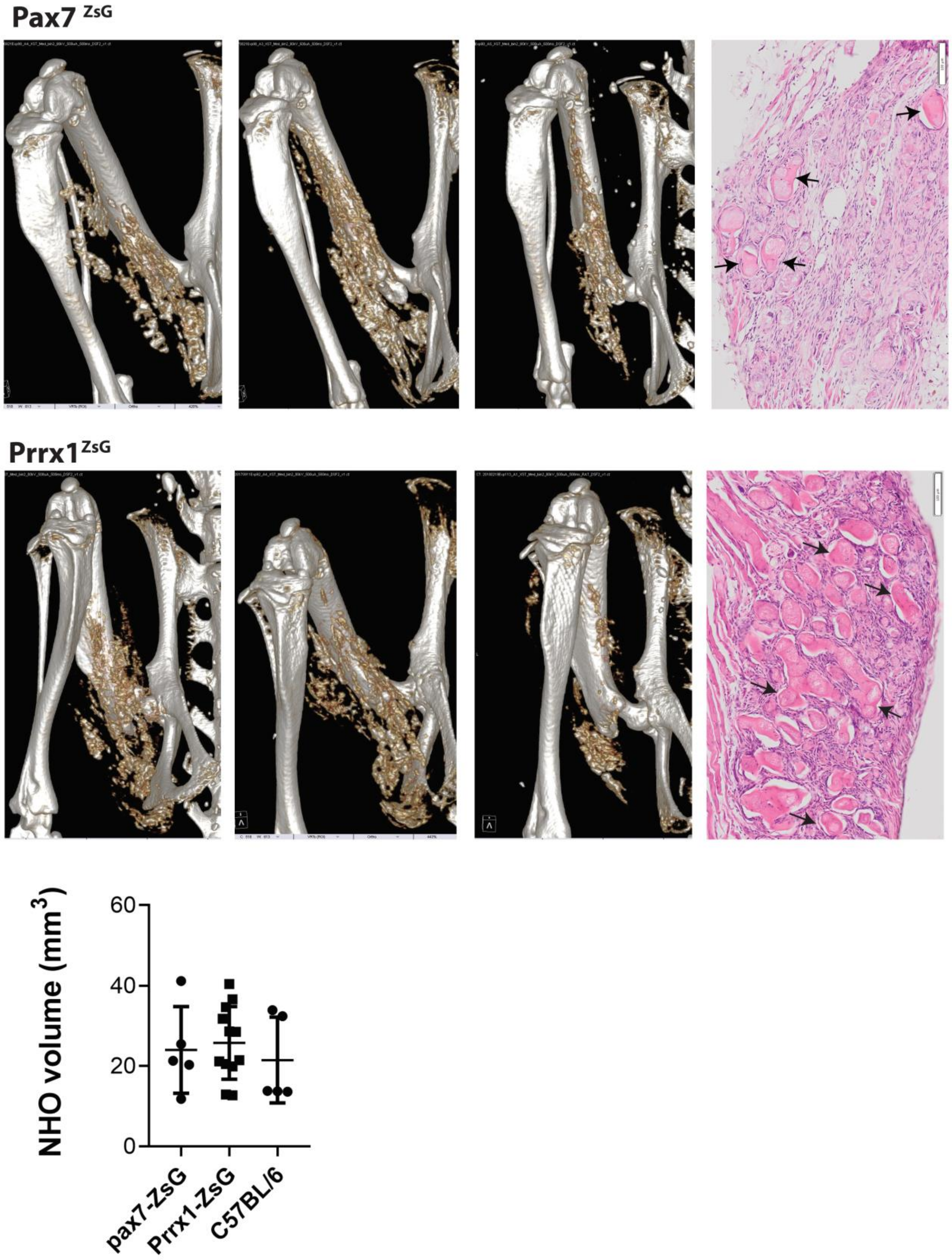


**Fig. S2.** *Pax7*ZsG and *Prrx1*ZsG mice underwent SCI surgery and i.m. injection of CDTX. NHO development was quantified by CT on day7 as representative images. Each point indicates one mouse and data was presented as mean ±S D. Statistic difference was calculated by Kruskal-Wallis test. Muscle were harvested on day28 and stained with Hematoxylin and eosin. Arrows indicates small bone nodules within necrotic muscle tissues.

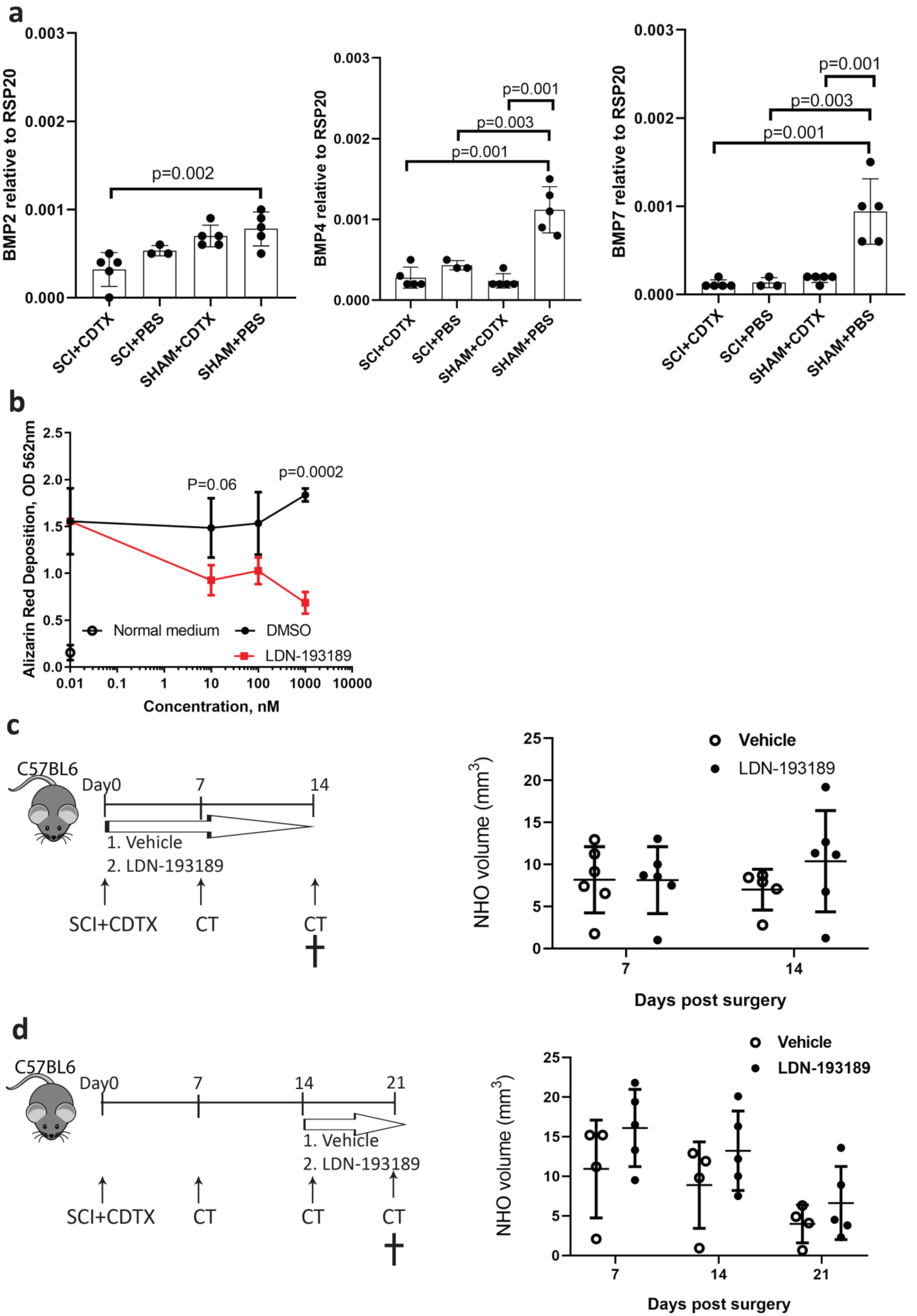
a
BMP2 relative to RSP20
BMP4 relative to RSP20
BMP7 relative to RSP20
p=0.002
p=0.001
p=0.003
p=0.001
p=0.001
p=0.003
p=0.001
SCI+CDTX
SCI+PBS
SHAM+CDTX
SHAM+PBS
b
Alizarin Red Deposition, OD 562nm
P=0.06
p=0.0002
Normal medium
DMSO
LDN-193189
Concentration, nM
c
C57BL6
Day0
7
14
1. Vehicle
2. LDN-193189
SCI+CDTX
CT
CT
NHO volume (mm3)
Vehicle
LDN-193189
Days post surgery
d
C57BL6
Day0
7
14
21
1. Vehicle
2. LDN-193189
SCI+CDTX
CT
CT
CT
NHO volume (mm3)
Vehicle
LDN-193189
Days post surgery

Figure S3

**Fig. S3.** BMP signalling is not required in the initiation and development of NHO. **a** *Bmp2*, *Bmp4* and *Bmp7* mRNA expression in muscles taken from SCI+CDTX, SCI+PBS, Sham+CDTX and Sham+PBS 4 days post-surgery were analysed by qRT-PCR. Expression was normalised by housekeeping gene *Rsp20*. Each point indicates one mouse and data was presented as mean±SD. Statistic difference was calculated by one-way ANOVA with Sidak multiple comparison analysis. **b** Mouse bone marrow mesenchymal cells were cultured with normal medium (open circles), osteogenic medium with recombinant BMP-2 (100ng/ml), DMSO vehicle (black full circles) or LDN-193189 (red squares) at indicated concentration. Mineralisation was quantified on day7 using Alizarin red and absorbance was read at 562nm. Data was presented as mean ± SD (triplicate/treatment) and statistic difference was analysed by two-way ANOVA and Bonferroni's multiple comparisons test. **c** Mice were treated with vehicle or LDN-193189 at 3 mg/kg twice daily by i.p. injection from day 0-14 and NHO were quantified by μCT on day7 and 14 (n=6 mice/group). **d** Mice were treated with vehicle or LDN-193189 (3 mg/kg twice daily) by i.p. injection from day 0-14 and NHO were quantified by μCT on day7, 14 and 21 (n=5 mice/group). Each point indicates one mouse and data was presented as mean ±S D. Statistic difference was calculated by Mann-Whitney test for comparison at each time point.

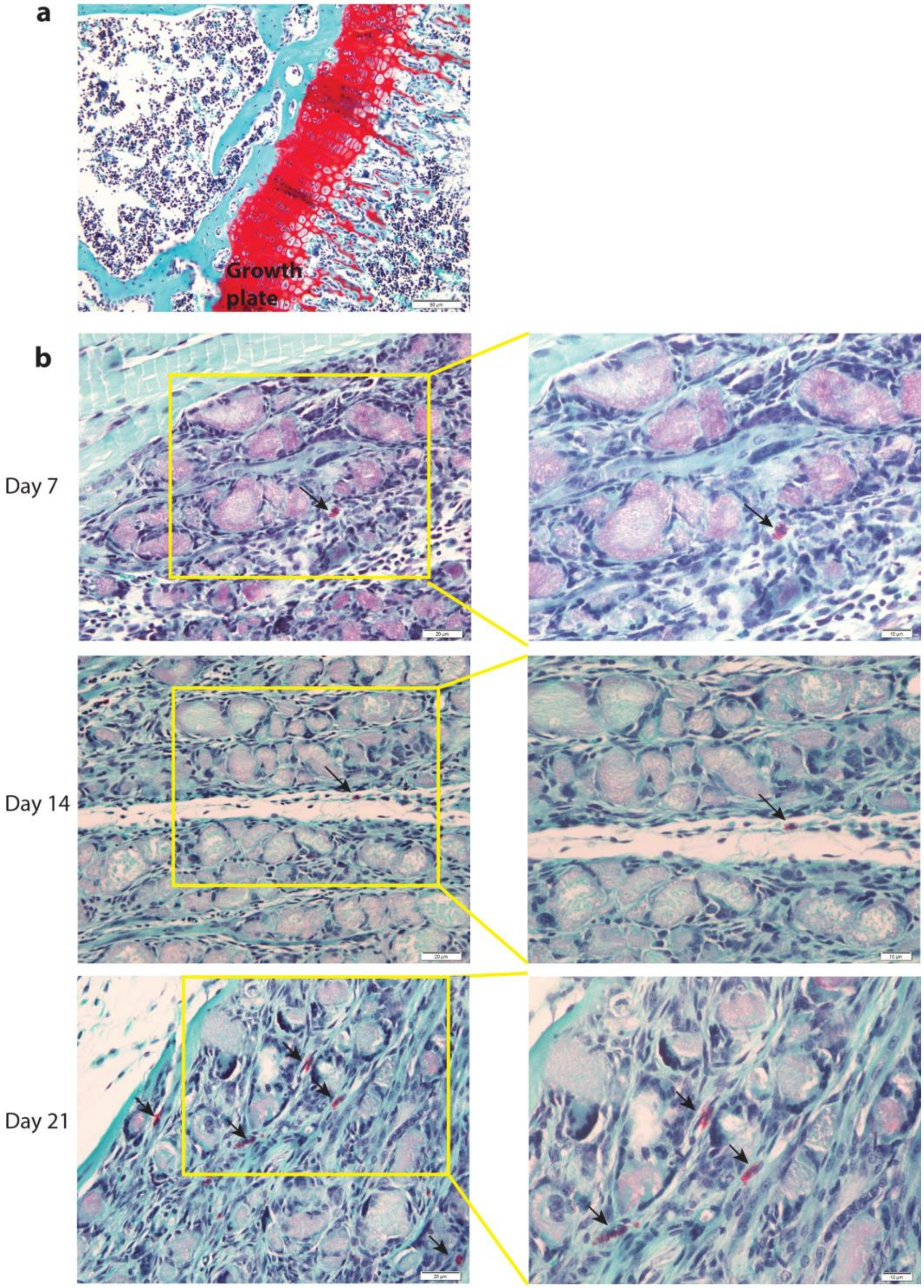


**Fig. S4.** NHO do not develop via endochondral ossification following SCI in mice. Safranin O/fast green staining was performed on injured muscle samples harvested 7, 14 and 21 days post SCI+CDTX surgery to examine whether NHO formation involves endochondral ossification. **a** Proteoglycan in growth plate stained bright red as positive control. **b** Occasional mast cells (arrows) with characteristic basophilic granules scattering in the injured muscle samples across day 7, 14 and 21 stained with Safranin O. However, bright red chondrocytes or cartilage matrix could not be detected in NHO at any time-point (n=3-6 mice per time point). Figures in the right column are higher magnification of yellow boxes in the left column. Scale bar: a 50 µm, b left =20µm, right =10µm.

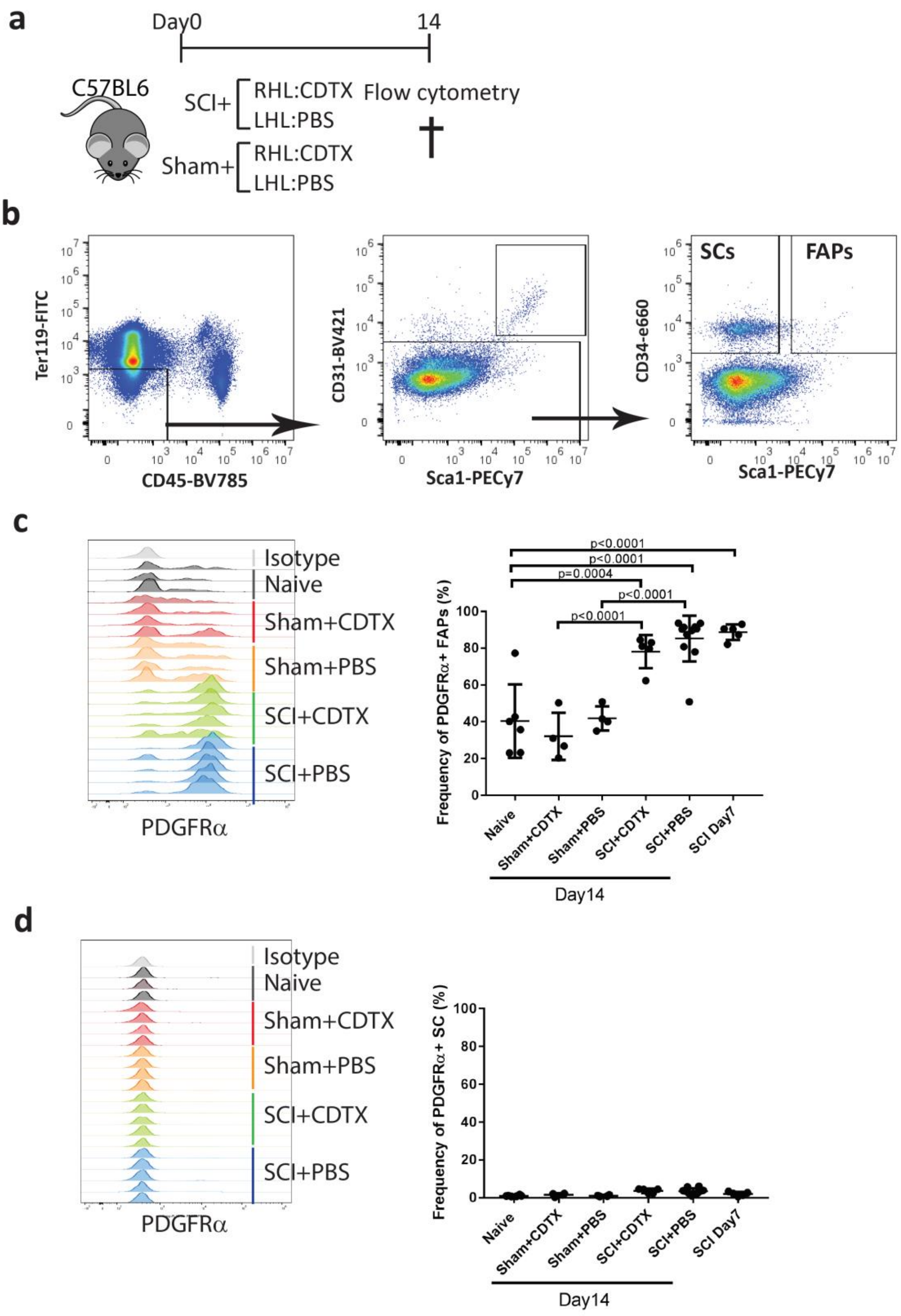


**Fig. S5.** SCI upregulates PDGFRα expression on FAPs but not SCs. **a** C57BL/6 mice received SCI or Sham surgery plus intramuscular injection of CDTX in RHL and PBS in LHL. Hamstring muscle cells were subsequently isolated 7 or 14 days post-surgery or from naïve mice. **b** FAPs and SCs were gated from live CD45-Ter119-CD31- cells and separated according to their expression of Sca1 and CD34. Overlayed histograms of PDGFRα expression on **c** $Sca1^+$ $CD34^+$ FAPs and **d** $Sca1^-$ $CD34^+$ SCs isolated from naïve (grey), sham + CDTX (red), sham + PBS (orange), SCI + CDTX (green), and SCI + PBS (blue) groups (n=3-5mice/group). Frequency of PDGFRα expression in FAPs and SCs of is shown as mean ± SD. Each dot represents a separate mouse. Significance was calculated by one-way ANOVA with Tukey's multiple comparison test.

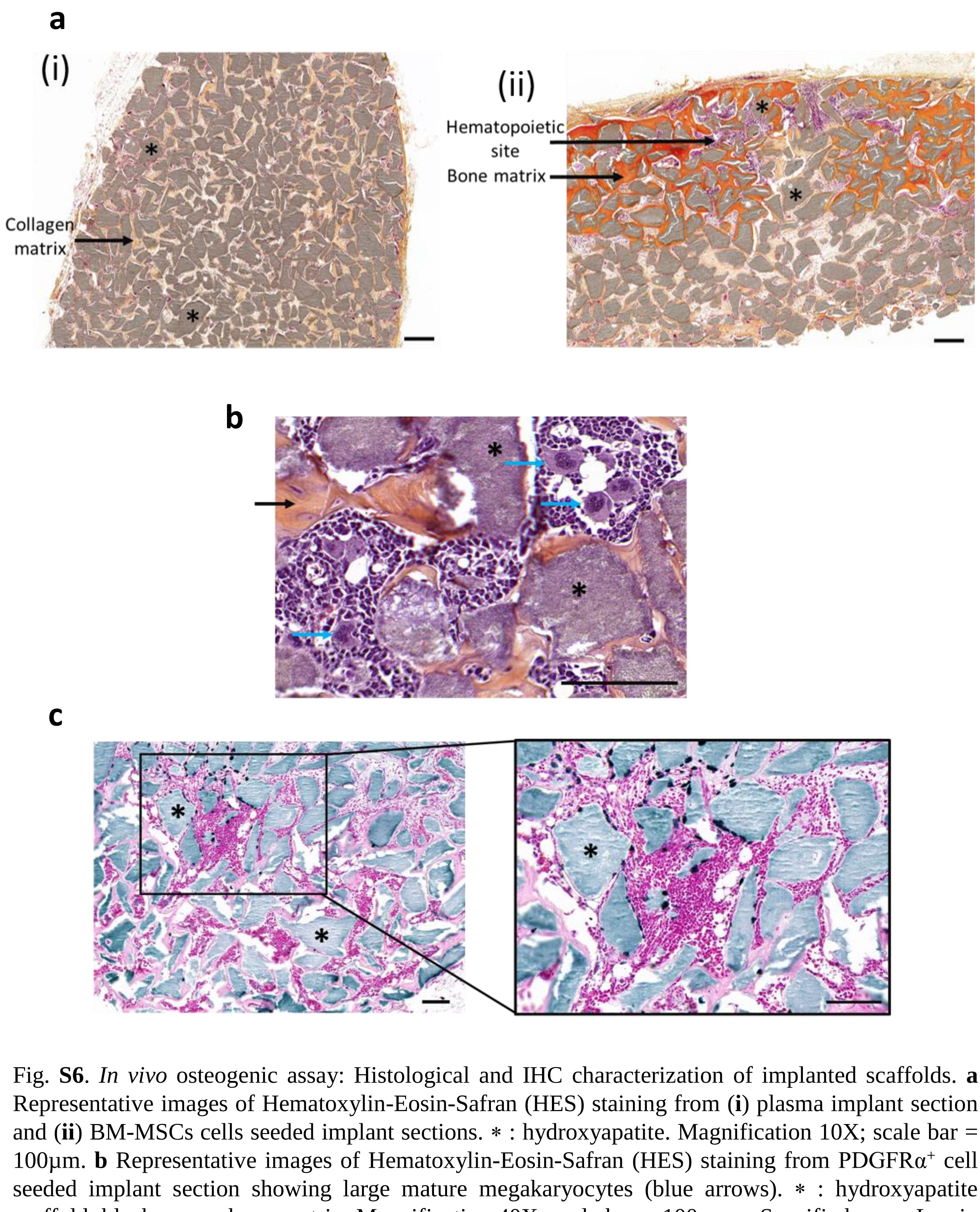


Fig. **S6**. *In vivo* osteogenic assay: Histological and IHC characterization of implanted scaffolds. **a** Representative images of Hematoxylin-Eosin-Safran (HES) staining from (**i**) plasma implant section and (**ii**) BM-MSCs cells seeded implant sections. ∗ : hydroxyapatite. Magnification 10X; scale bar = 100µm. **b** Representative images of Hematoxylin-Eosin-Safran (HES) staining from PDGFR$\alpha^+$ cell seeded implant section showing large mature megakaryocytes (blue arrows). ∗ : hydroxyapatite scaffold; black arrow: bone matrix. Magnification 40X; scale bar = 100µm. **c** Specific human Lamin A/C staining of a representative PDGFR$\alpha^+$ cell seeded implant section. ∗ : hydroxyapatite scaffold. Magnification 10X and 20X; scale bar = 100µm.
