## Supplementary figures for "Spinal cord injury reprograms muscle fibro-adipogenic progenitors to form heterotopic bones within muscles"

**Supplementary Table 1**. Percentage of phenotypic SCs, FAPs and endothelial cells in non-hematopoietic cells from the hamstring muscles of *Pax7*$^{ZsG}$ and *Prrx1*$^{ZsG}$ mice at 14 days post PBS or CDTX injection measured by flow cytometry.

| Mouse strains | Intramuscular injections | Satellite cells | Fibro-adipogenic progenitors | Endothelial cells |
|---|---|---|---|---|
| *Pax7*$^{ZsG}$ | PBS | 11.0 ± 2.6 | 15.7 ± 5.1 | 0.56 ± 0.12 |
| *Prrx1*$^{ZsG}$ | PBS | 11.1 ± 2.3 | 11.8 ± 1.8 | 0.66 ± 0.19 |
| *Pax7*$^{ZsG}$ | CDTX | 9.9 ± 0.5 | 16.72 ± 7.3 | 0.72 ± 0.33 |
| *Prrx1*$^{ZsG}$ | CDTX | 9.8 ± 0.9 | 15.2 ± 3.4 | 0.81 ± 0.23 |

Data are presented as mean ± SD (n=4 mice/group for *Pax7*$^{ZsG}$ and n=3 mice/group for *Prrx1*$^{ZsG}$). There was no significant difference between the different strains as determined by one-way ANOVA.

**Supplementary Table 2.** Genotyping PCR reagents and conditions

| Genes | Primers | Primer sequences | PCR Conditions |
|---|---|---|---|
| Pax7-CreERT2 | Pax7 common F | 5'GCTGCTGTTGATTACCTGGC | 94 °C 15s, 60°C 30s, 72°C 60s |
| | Pax7 wt R | 5'CTGCACTGAGACAGGACCG | |
| | Pax7 Mutant R | 5'CAAAAGACGGCAATATGGTG | |
| Rosa26R-Zsgreen | Rosa26 wt F | 5' AAG GGA GCT GCA GTG GAG TA | 94 °C 15s, 62°C 30s, 72°C 60s |
| | Rosa26 wt R | 5' CCG AAA ATC TGT GGG AAG TC | |
| | Rosa26-Zsgreen Mut R | 5' GGC ATT AAA GCA GCG TAT CC | |
| | Rosa26-Zsgreen Mut F | 5' AAC CAG AAG TGG CAC CTG AC | |
| Prrx1Cre | Cre forward | 5' GAGTGATGAGGTTCGCAAGA | 94 °C 15s, 58°C 30s, 72°C 60s |
| | Cre reverse | 5' CTACACCAGAGACGGAAATC | |

**Supplementary Table 3**. Antibodies used in the study.

1) **Flowcytometry (anti-mouse)**

| Anti-mouse | Catalogue | Company | Clone | Dilution |
|---|---|---|---|---|
| B220- Biotin | 103204 | Biolegend | RA3-6B2 | 1:200 |
| CD3-Biotin | 100304 | Biolegend | 145-2C11 | 1:200 |
| CD11b- Biotin | 101204 | Biolegend | M1/70 | 1:200 |
| CD31-BV421 | 102423 | Biolegend | 390 | 1:200 |
| CD34-e660 | 50-0341-82 | eBioscience | RAM34 | 1:75 |
| CD45-BV785 | 103149 | Biolegend | 30-F11 | 1:200 |
| F4/80-PE | 123110 | Biolegend | BM8 | 1:500 |
| GR1- Biotin | 108404 | Biolegend | RB6-8C5 | 1:200 |
| α7integrin-PE | 53-0010-05 | Ablab | R2F2 | 1:3000 |
| PDGFRα (CD140a)-BB700 | 742176 | BD Bioscience | APA5 | 1:200 |
| PDGFRα (CD140a)-PE | 135905 | Biolegend | APA5 | 1:100 |
| Sca1-PECy7 | 108114 | Biolegend | D7 | 1:300 |
| Ter119- Biotin | 116204 | Biolegend | TER-119 | 1:200 |
| Ter119-FITC | 116206 | Biolegend | TER-119 | 1:300 |
| Fixable Viability Stain 700 | 564997 | BD Bioscience | - | 1:15000 |
| SAV-APCCy7 | 554063 | BD Bioscience | - | 1:200 |
| B220-PercpCy5.5 | 103236 | Biolegend | RA3-0B2 | 1:200 |
| Ter119-PercpCy5.5 | 116228 | Biolegend | TER-119 | 1:200 |
| CD3-PercpCy5.5 | 100328 | Biolegend | 145-2C11 | 1:200 |
| Ly6G-APCCy7 | 127624 | Biolegend | 1A8 | 1:200 |
| CD11b-BV510 | 101263 | Biolegend | M1/70 | 1:200 |
| CD3-Pacific blue | 100334 | Biolegend | 145-2C11 | 1:200 |
| GR1-Pacific blue | 108430 | Biolegend | RB6-8C5 | 1:200 |
| B220-Pacific blue | 103227 | Biolegend | RA36B2 | 1:200 |
| CD11b-Pacific blue | 101224 | Biolegend | M1/70 | 1:200 |
| Ter119-Pacific blue | 116231 | Biolegend | TER-119 | 1:200 |
| CD5-Pacific blue | 100642 | Biolegend | 53-7.3 | 1:200 |

| Anti-mouse | Catalogue | Company | Clone | Dilution |
|---|---|---|---|---|
| CD31-PE | 102407 | Biolegend | 390 | 1:200 |
| FITC BrdU Flow Kit | 559619 | BD Biosciences | | 1:20 |
| Annexin V-FITC | 640906 | Biolegend | | 1:50 |

### 2) Flowcytometry (anti-Human)

| Antibody | Catalogue | Company | Clone | dilution |
|---|---|---|---|---|
| huCD31-PE | 555446 | BD Biosciences | WM59 | 1:10 |
| huCD34-APC | IM2472 | Beckman Coulter | 581 | 1:10 |
| huCD45-APC | IM2473 | Beckman Coulter | J33 | 1:10 |
| huCD73-PE | 550257 | BD Biosciences | AD2 | 1:10 |
| huCD90-APC | 559869 | BD Biosciences | 5E10 | 1:10 |
| huCD105-PE | PN A07414 | Beckman Coulter | 1G2 | 1:10 |
| huCD56-PE | 555516 | BD Biosciences | B159 | 1:10 |
| huPDGFRα-Biotin | BAF1322 | R&D Systems | - | 1:10 |
| Streptavidin APC/Cy7 | 2626040 | Sony | - | 1:200 |
| 7-AAD (7-Aminoactinomycin D) | A1310 | Molecular probes | - | |
| huLamin A/C | ab108595 | Abcam | EPR4100 | 1:100 |
| hu-muOsterix/SP7 | ab22552 | Abcam | - | 1:100 |

### 3) Immunofluroescence staining

| Antibody | Catalogue | Company | Concentration |
|---|---|---|---|
| Collagen Type I | C7510-13 | US Biological | 1 ug/ml |
| Osteocalcin | ALX-210-333 | EnzoLife Sciences | 1ug/ml |
| Rabbit IgG control | 31235 | ThermoFisher | 1ug/ml |
| biotin-labelled goat-anti-Rabbit IgG secondary antibody | BA-1000 | Vector Labs | |

| | | |
|---|---|---|
| Streptavidin, Alexa Fluor™ 647 conjugate | S21374 | Invitrogen |

4) **Western blot**

| Antibody | Catalogue | Company | Clone | Dilution |
|---|---|---|---|---|
| p-Akt (S473) | 4060s | Cell Signaling | | 1:1000 |
| Total Akt (c67E7) | 4691 | Cell Signaling | | 1:1000 |
| IRDye® 800CW Donkey anti-Rabbit IgG (H + L) | LCR -926-32213 | Licor | | 1:15000 |

**Supplementary Table 4: qRT-PCR primer probe sets**

| | Catalogue | Company |
|---|---|---|
| SensiFast | BIO65054 | Bioline |
| TaqMan™ Fast Advanced Master Mix | 4444557 | ThermoFisher |
| Rsp20 gene expression assay | Mm02342828_g1 | ThermoFisher |
| Bmp2 gene expression assay | Mm01340178_m1 | ThermoFisher |
| Bmp4 gene expression assay | Mm00432087_m1 | ThermoFisher |
| Bmp7 gene expression assay | Mm00432102_m1 | ThermoFisher |

**Supplementary Table 5.** Other reagents.

| Reagents | Company | Catalogue number |
|---|---|---|
| Tamoxifen | Signa-Aldrich | T5648 |
| Cardiotoxin | Latoxan | L8102 |
| Skeletal muscle dissociation kit | Miltenyi Biotec | 130-098-305 |
| Recombinant Mouse PDGF-BB (carrier-free) | Biolegend | RUO-558802 |
| Recombinant human BMP2 | Peprotech | 120-02C-10 |
| LDN-193189 | Cayman Chemical | 19396 |
| Dasastinib | Tocris | 6793 |
| DAPI | Signa-Aldrich | D5942 |
| ProLong™ Gold Antifade Mountant | Invitrogen | P36930 |
| Tissue-Tek® O.C.T.™ Compound | Tissue-Tek | IA018 |
| Superfrost Plus™ Adhesion Microscope Slides | Thermo Fisher Scientific | MENSF41296SP |
| Mx 35 premier + blades | Thermo Scientific | 3052835 |
| cOmplete™ ULTRA Tablets, EDTA-free, glass vials Protease Inhibitor Cocktail | Roche | 05892953001 |
| 4-12% Bis-Tris pre-cast gel | Invitrogen | NW04122 |
| Thermofisher nitrocellulose mini stacks | ThermoFisher | B23002 |
| Odyssey Blocking Buffer | Licor | 921-50000 |
| Restore PLUS Western Blot Stripping Buffer | ThermoFisher | 46430 |
| Paraformaldehyde | Signa-Aldrich | P6148 |
| EDTA disodium salt | Astral Scientific | BIOEB0185-500g |
| Alizeran red S | Signa-Aldrich | A5533 |
| Cetylpyridinium chloride | Signa-Aldrich | C0732 |
| β-Glycerophosphate disodium salt hydrate | Signa-Aldrich | A5533 |
| Dexamethasone | Hospira | 433449 |
